# DMLDA-LocLIFT: Identification of multi-label protein subcellular localization using DMLDA dimensionality reduction and LIFT classifier

**DOI:** 10.1101/2020.03.06.980441

**Authors:** Qi Zhang, Shan Li, Bin Yu, Qingmei Zhang, Yan Zhang, Qin Ma

**Affiliations:** College of Mathematics and Physics, Qingdao University of Science and Technology, Qingdao 266061, China; Artificial Intelligence and Biomedical Big Data Research Center, Qingdao University of Science and Technology, Qingdao 266061, China; School of Mathematics and Statistics, Central South University, Changsha 410083, China; School of Life Sciences, University of Science and Technology of China, Hefei 230027, China; College of Electromechanical Engineering, Qingdao University of Science and Technology, Qingdao, 266061, China; Department of Biomedical Informatics, College of Medicine, The Ohio State University, Columbus, Ohio 43210, USA

**Keywords:** Subcellular localization, Multi-information fusion, Direct multi-label linear discriminant analysis, Multi-label learning with label-specific features

## Abstract

**Background:** Multi-label proteins occur in two or more subcellular locations, which play a vital part in cell development and metabolism. Prediction and analysis of multi-label subcellular localization (SCL) can present new angle with drug target identification and new drug design. However, the prediction of multi-label protein SCL using biological experiments is expensive and labor-intensive. Therefore, predicting large-scale SCL with machine learning methods has turned into a hot study topic in bioinformatics.

**Methods:** In this study, a novel multi-label learning means for protein SCL prediction, called DMLDA-LocLIFT, is proposed. Firstly, the dipeptide composition, encoding based on grouped weight, pseudo amino acid composition, gene ontology and pseudo position specific scoring matrix are employed to encode subcellular protein sequences. Then, direct multi-label linear discriminant analysis (DMLDA) is used to reduce the dimension of the fused feature vector. Lastly, the optimal feature vectors are input into the multi-label learning with Label-specIfic FeaTures (LIFT) classifier to predict the location of multi-label proteins.

**Results:** The jackknife test showed that the overall actual accuracy on Gram-negative bacteria, Gram-positive bacteria, and plant datasets are 98.60%, 99.60%, and 97.90% respectively, which are obviously better than other state-of-the-art prediction methods.

**Conclusion:** The proposed model can effectively predict SCL of multi-label proteins and provide references for experimental identification of SCL. The source codes and data are publicly available at https://github.com/QUST-AIBBDRC/DMLDA-LocLIFT/.

## 1. Introduction

Cell growth, development, reproduction, and other vital activities are inextricable with the action of proteins. In the level of cell, proteins only fulfill a function in particular subcellular locations [1]. These positions offer a certain chemical environment and a series of interacting pairs [2, 3], enabling proteins to perform their functions correctly [4]. SCL is critical for protein function. Abnormal proteins are tightly related to human diseases, such as brucellosis [5], primary human liver tumors [6], breast cancer [7], preeclampsia [8], etc. Moreover, understanding the location of proteins can present new angle with drug design and drug target identification [9].

As we can know, study has got into the post-genomic era, the quantity of protein sequences in the database has gradually increased. It is far from satisfying the research which needs to search for the subcellular position of proteins using experimental methods. So how to predict protein SCL with machine learning becomes a question that should be solved quickly. In the past two decades, several prediction methods [10, 11] had been proposed to solve this problem, but these prediction approaches were developed based on a single labeling system in which every protein only had one subcellular position [12]. However, the experiment shows that more and more proteins can occur in two or more different subcellular locations simultaneously [13]. These multi-label proteins play a more important role in cell vital movements, so the learning of SCL of a multi-label protein is particularly important.

Predicting multi-label SCL through machine learning can not only save man power, material resources, and financial resources but also provide some references for experimental methods. To build a SCL model by the method of calculation, the foremost task is to draw the feature information from protein sequences. The commonly used feature extraction methods are mainly derived from structural information, physicochemical properties, evolutionary information, and sequence information. To reveal the sequence information of animal proteins, Cheng et al. [14] used pseudo amino acid composition (PseAAC) to draw the useful information from amino acid residues of animal protein. Huang et al. [15] encoded protein sequences by integrating protein sequences information, physicochemical information and evolutionary information. Wang et al. [16] improved the position specific scoring matrix with segmented amino acid composition to extract characteristic message from Gram-positive bacteria and Gram-negative bacteria. Shen and Chou [17] had proposed a new method Gneg-mPLoc for identifying gram-negative bacteria, which used three coding methods gene ontology, function domain, and sequence evolution to extract features from Gram-negative bacteria.

Feature fusion can generate high-dimensional matrix, which includes lots of superfluous information, and may influence the performance of the classifier seriously. Dimensional reduction is able to assist us reduce noise and eliminate superfluous information. It is also widely applied in pattern recognition and classification. Today, researchers have introduced multiple dimensionality reduction means in the study of SCL prediction, including principal component analysis (PCA) [18], multi-label informed latent semantic indexing (MLSI) [19], multi-label dimensionality reduction via dependency maximization (MDDM) [20], multi-label feature extraction algorithm via maximizing feature variance and feature-label dependence (MVMD) [21] and etc. Zhang et al. [22] proposed manifold regularized discriminant feature selection (MDFS), which utilized label correlation to establish an optimization framework and used iterative optimization algorithm to solve convexity optimization problems, thus greatly improving the performance of prediction. Zhang and Zhou [20] proposed a dimensionality reduction method MDDM, which used projection strategy to project raw data into lower-dimensional feature space and can make full use of the reliance between related class labels and primitive features. Lin et al. [23] proposed a max-dependency and min-redundancy (MDMR) method, which combined mutual information with maximum correlation and minimum redundancy to solve the multi-label dimension explosion issue and select the best subset for multi-label learning.

Apart from the effect of dimensionality reduction methods, the selection of the classifier is also critical to the prediction. The commonly used multi-label classification methods include multi-label k-nearest neighbor (ML-KNN) [24], multi-label neural network (BP-MLL) [25], multi-label learning by instance differentiation (InsDif) [26], multi-label learning by exploiting label correlations local (ML-LOC) [27] and multi-label learning with label-specific features (LIFT) [28] etc. Wang et al.[29] used the classifier chain (ECC) classifier set to predict the multi-label proteins of Gram-negative bacteria and Gram-positive bacteria. After testing, the overall actual accuracy (OAA) and overall location accuracy (OLA) of the two datasets were 92.4%, 94.1% and 94.0%, 94.4%, respectively. Wan et al. [30] used a multi-label support vector machine classifier through selecting virus proteins and plant proteins as the benchmark datasets. Jackknife test showed the overall actual accuracy (OAA) and overall location accuracy (OLA) of the two datasets were 93.7%, 97.2% and 93.6%, 95.6%, respectively. Wan et al. [31] used Multi-label LASSO as a classifier to predict the human proteins dataset and used the LOOCV to assess the performance of the model. The overall actual accuracy (OAA) and overall location accuracy (OLA) were 74.8% and 84.6%, respectively, etc.

Although some achievements have been made in protein SCL prediction using machine learning means, there are still many aspects for melioration. Firstly, in the process of multi-label learning, the characteristic information of protein sequences characterizing SCL has not been fully elucidated. Then, high dimensional matrix after fusion will produce many redundant features. High-dimensional data will not only reduce the speed of operation but also affect the results of the model prediction. It is an urgent problem to be solved how to select a suitable dimensionality reduction method to eliminate noise information, and then reduce the running time of the model and improve the prediction performance. Finally, the experimental data of the multi-label protein subcellular shows geometric growth. It is essential to design a precise, fast and effective prediction tool for SCL.

Enlightened by this, we put forward a new protein SCL prediction method called DMLDA-LocLIFT based on multi-label learning. Firstly, Extracting information from protein sequences by pseudo amino acid composition (PseAAC), dipeptide composition (DC), encoding based on grouped weight (EBGW), gene ontology (GO) and pseudo position-specific scoring matrix (PsePSSM), and the best parameter *λ* values, *ξ* values, and *L* values of the model are determined by jackknife test. Secondly, the feature extraction information are combined as the initial features. Compared with other dimensionality reduction methods, DMLDA can capture the effective feature representation by removing the redundancy from raw feature space. At last, the first-best feature subsets are input into the LIFT classifier to predict the location of multi-label protein SCL, which achieves better prediction accuracy than ML-LOC, ML-KNN and INSDIF classifiers. The results indicate that the proposed DMLDA-LocLIFT method can significantly increase the prediction accuracy of protein SCL.

## 2. Materials and methods

### 2.1. Datasets

In the light of Chou’s five-step rule [32], using strict benchmark dataset is the first step for establishing reliable models. Benchmark datasets are generally composed of training dataset and test dataset. The former is to train the model, and latter is to assess the model. In this paper, three different multi-label protein subcellular datasets are selected, namely, Gram-negative bacteria dataset [33], Gram-positive bacteria dataset [33] and plant dataset [30]. The Gram-negative bacteria dataset has 1456 locative proteins in the dataset. It has 1392 distinct protein sequences located in 8 subcellular positions, of which 1328 belong to one subcellular position, 64 belong to two positions and none to three or more positions. The Gram-positive bacteria dataset has 523 locative proteins in the dataset. It includes 519 distinct protein sequences located in 4 subcellular positions, where 515 belong to one subcellular position, 4 belong to two positions and none to three or more positions. The subcellular location name and number of the Gram-negative bacteria and Gram-positive bacteria datasets are shown in Supplementary Table S1 and S2. The plant dataset is used as an independent test set to verify the validation of the model. There are 1055 locative proteins in the dataset. The dataset has 978 distinct protein sequences located in 12 subcellular positions, of which 904 belong to one subcellular position, 71 belong to two locations, 3 belong to three positions and none to four or more positions. The specific distribution is shown in Table 1. The homology of protein sequences in the three datasets are limited to below 25%.

**Table 1.**
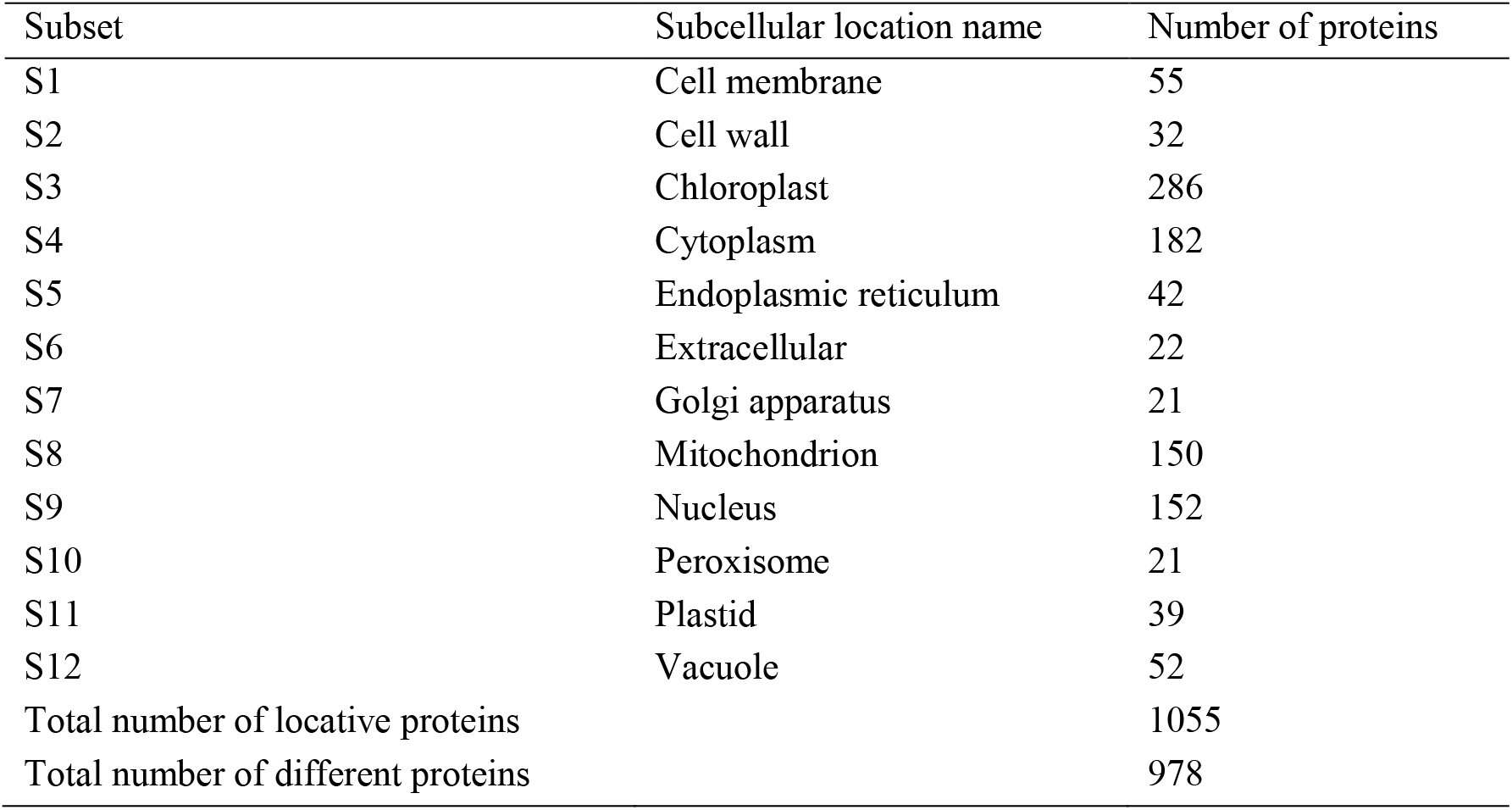
Breakdown of the plant bacterial benchmark dataset.

### 2.2. Feature extraction

So as to predict the location of multi-label protein subcellular, the first task is to draw the feature from protein sequence and transform the protein sequence information into digital information. In this article, we use five feature extractions to obtain the information of protein sequence, namely PseAAC, PsePSSM, EBGW, DC and GO.

#### 2.2.1. Pseudo Amino Acid Composition

PseAAC [34–36] mainly uses sequence information and physical and chemical properties in protein sequences for feature extraction. Currently, the researchers have widely used the method of pseudo-amino acid composition in proteomics. The means maps the protein sequence into the following feature vectors:

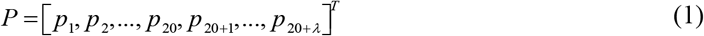

Every component is defined as:

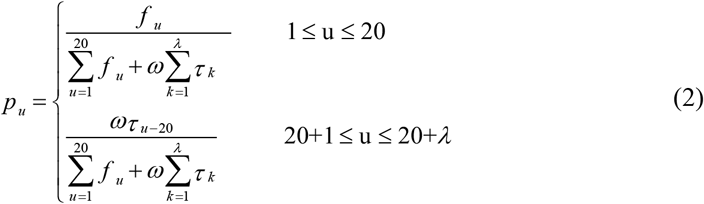

among them, *ω* is the weight factor and the value is 0.05 in the literature [37]. *τ*_*k*_ is the *k* nearest correlation factor, which represents sequence information between each amino acid. *f*_*u*_ represents the frequency of the *u* -th amino acid. Therefore, the former 20 dimensions of the eigenvector *P* are the composition of amino acids, the other *λ* dimensions are the correlation factors of different levels reflecting the sequence information of amino acids. Because the three standard datasets selected in this paper require that the length of protein sequence should not be too short, the smallest sequence length is 50, so the selection of *λ* is *λ* ≤ 50. In this paper, by choosing different parameter *λ*, the first-best *λ* can be determined by the precision of the prediction results.

#### 2.2.2. Pseudo position-specific scoring matrix

In this paper, PSI-BLAST served to obtain PSSM of protein sequence. First, download a non-redundant protein database provided by NCBI. Secondly, all protein sequences are compared with this non-redundant database by using the PSI-BLAST program. In this course, set the threshold E to 0.001, and the maximum number of iterations is 3. BLOSUM62 is selected as the contrast matrix. The other parameters take the default value, and the position-specific score matrix (PSSM) of every sequence is obtained. Drawing features from protein sequence with *L* length, it is obtained that its corresponding position-specific score matrix (PSSM) [38] is *L*×20 matrix, such as formula (3).

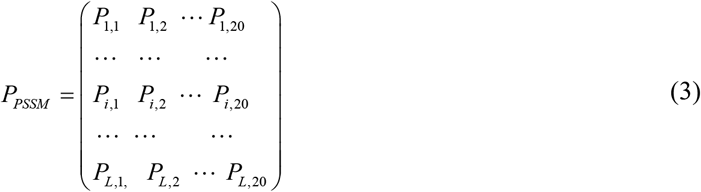

Firstly, the above matrix is standardized, and the value of PSSM matrix is transformed into 0 to 1 by the formula (4):

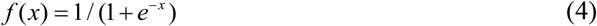

In this article, PSePSSM algorithm [39, 40] is applied to draw the features from protein sequences. And process of feature extraction is as follows:

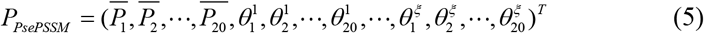

among them, 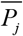 denotes the component of amino acid *j* in PSSM. 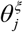 represents the *ξ* -order relevant factors about amino acid type *j*. As can be seen from formula (5), a sequence can be produced a 20 + 20×*ξ* -dimensional feature vector.

#### 2.2.3. Encoding based on grouped weight

Zhang [41] et al. starting from the physicochemical properties about amino acids, used the idea of “coarse-grained” and “grouping” to reduce protein sequences into binary feature sequences, and introduced the normal weight function based on the feature sequences to realize the Encoding based on grouped weight (EBGW) [42, 43] of protein sequences. On the basis of physical and chemical properties (hydrophobicity and charge) of the 20 basic amino acids which constitute the protein sequence, they were divided into four categories:

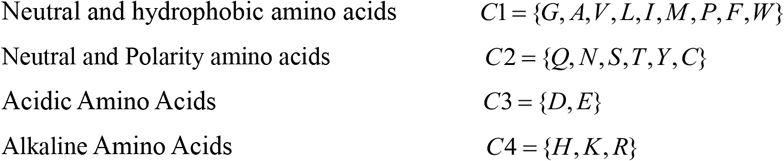

Combining *C*1, *C*2, *C*3, *C*4 in pairs and obtain {*C*1, *C*2} vs {*C*3,*C*4}, {*C*1, *C*3} vs {*C*2,*C*4} and {*C*1, *C*4} vs {*C*2,*C*3} three new divisions. Let *P* = *p*_1_*p*_2_ ⋯ *p*_*N*_ be a protein sequence, We can use three homomorphic maps *f*_1_(*P*), *f*_2_(*P*), *f*_3_(*P*) to transform *P* into three binary sequences. The definition is as follows:

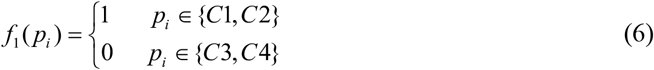

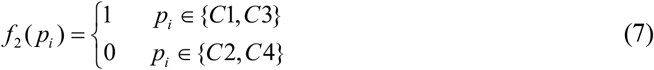

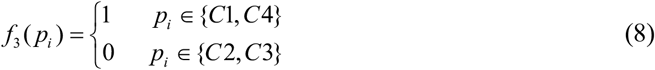

*f*_1_(*P*), *f*_2_(*P*), *f*_3_(*P*) is a binary sequence with three lengths of *n*. These sequences are divided into several subsequences with increasing lengths in turn. Setting a fixed parameter *L*, the subsequence can be represented as ⌊*kn* / *L*⌋, *k* = 1, 2,⋯*L*, each *f*_*i*_(*P*) generates an *L* -dimensional feature vector. Therefore, for any sequence of length *L*, a 3*L* -dimension vector can be produced.

#### 2.2.4. Dipeptide composition

The dipeptide composition (DC) model [44] is to calculate the frequency of occurrence about amino acid pairs. The vector elements extracted is shown in equation (9):

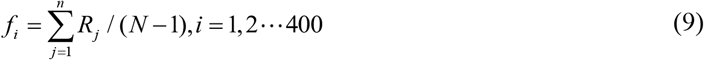

where *P*_*i*_ denotes the frequency of amino acid pairs, *N* represents the length of sequence. There are 20 × 20 = 400 ways to combine 20 amino acid forms into amino acid pairs.

#### 2.2.5. Gene ontology

For a protein sequence *P*_*i*_ in the dataset, BLASTP (http://blast.ncbi.nlm.nih.gov/Blast.cgi) is used to search for *P*_*i*_ homologous proteins in SWISS-PROT database. And parameter E is set to 0.001. In the obtained homologous sequence, protein sequences of similarity ≥ 60% is screened out and formed a protein set with *P*_*i*_, which is recorded as *S*_*i*_. For each protein sequence in set *P*_*i*_, the GO information [45, 46] of the corresponding protein is searched in the GO database by using the index number of the sequence.

Equation (10) is used to construct a GO binary vector. The elements in the vector indicate whether a GO number exists or not. Due to many GO numbers, delete those GO numbers that do not exist in the corresponding dataset to reduce redundant information.

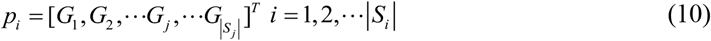

where |*S*_*i*_| is the size of the *S*_*i*_ set, 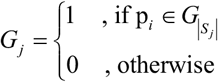.

### 2.3. Direct multi-label discriminant analysis (DMLDA)

A multi-label dataset containing *n* samples and *K* labels are defined as 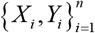, where *X*_*i*_ ∊ *R*^*d*^, *Y*_*i*_ ∊{0,1}^*K*^. If *X*_*i*_ pertain to the *k-th* label, then *Y*_*i*_(*k*) = 1. Each label is treated as a category, and the samples are divided into *K* groups. The number of samples in the category *k* is represented by *n*_*k*_. If *Y*_*i*_ ∊{0,1}^*K*^ is the label indicator vector of *X*_*i*_ and *y*_(*k*)_∊{0,1}^*n*^ is the label indicator vector of category *k*, then the sample set is *X* = [*X*_1_,⋯*X*_*n*_] and the label indicator set is *Y* = [*Y*_1_,⋯*Y*_*n*_]^*T*^ = [*y*_(1)_,⋯*y*_(*K*)_].

Considering that labels are interrelated, the tag relationship between two classes in the DMLDA algorithm [47] is defined as follows.

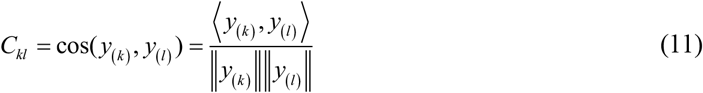

Compared with the single-label dataset, the samples containing multiple labels are computed repeatedly. Let *Z* = [*z*_1_,⋯*z*_*n*_]^*T*^, correct the repeated computation problem by normalizing the matrix.

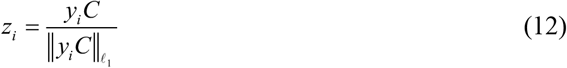

Computing between-class scatter matrices:

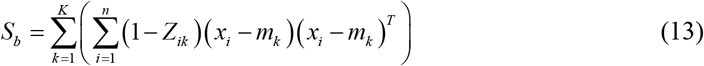

Computing within-class scatter matrices:

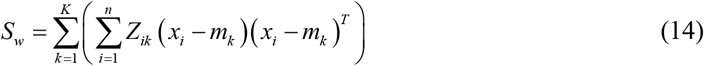

among them, *m*_*k*_ is the average value of the *k-th* class and *m* is the global average value of the multi-tagged dataset.

Solving the objective function of MLDA:

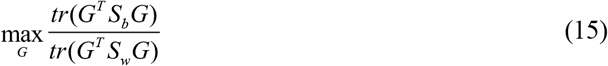

where *G* is a transformation matrix projecting dataset into low-dimensional space, and the matrix after dimensionality reduction becomes: *T* = *G*′*X*.

### 2.4. Multi-label learning with Label-specIfic FeaTures (LIFT)

Zhang et al. [28] studied a strategy of learning from a multi-label dataset, which utilizes the specific features of labels and is conducive to the recognition of different types of labels. Therefore, a credible algorithm called LIFT is proposed, i.e., multi-label learning with label specific features. At first, LIFT constructs the unique features of every tag by clustering the instances [48], and then each classifier is induced from the generated features for prediction.

Suppose that given *m* multi-label training sample *D* = {(*x*_*i*_,*Y*_*i*_) | 1 ≤ *i* ≤ *m*}, where *x*_*i*_ ∊ *X* is the feature vector of *d* -dimensional and *Y*_*i*_ ∊ *Y* is the corresponding category label of *x*_*i*_. The LIFT classification algorithm completes the construction of tag features and the induction of classification model through the following specific steps.

Firstly, the sample data of each category are clustered, and the clustering centers of positive and negative examples in each category are obtained. In this way, the features of each tag feature can be captured effectively, thus providing appropriate distinguishing information to facilitate its recognition process. Specifically, for the class label *l*_*k*_ ∊*Y*, positive training instance set *P*_*k*_ and negative training instance set *N*_*k*_ are represented via equation (16) and equation (17), respectively.

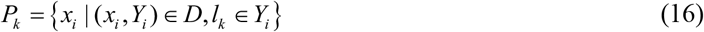

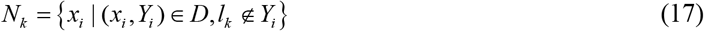

where, *P*_*k*_ and *N*_*k*_ contain training examples with and without *l*_*k*_ label, respectively.

Secondly, in order to further study the properties of *P*_*k*_ and *N*_*k*_, LIFT adopts clustering method. In this paper, *K* -means clustering [49] is applied to positive training instance set *P*_*k*_ and negative training instance set *N*_*k*_. Assuming that *P*_*k*_ is divided into 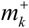 disjoint clusters and *N*_*k*_ is divided into 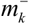 disjoint clusters, and then two groups of clustering centers 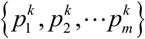 and 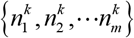 can be obtained respectively. Because multi-label learning may also have an imbalance problem, the negative training instance of each class label is much larger than that of the positive training instance, that is, |*P*_*k*_|≤|*N*_*k*_|. To avoid the disturbance caused by imbalance, LIFT sets *P*_*k*_ and *N*_*k*_ to the same number of clusters, namely 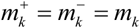. Therefore, the information obtained from positive training examples and negative training examples can be treated equally. Specifically, the number of clusters divided by *P*_*k*_ and *N*_*k*_ is set as equation (18):

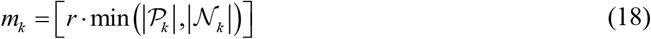

where | · | represents the cardinality set back, *r* ∊[0,1] is the ratio parameter of the number of reserved clusters.

By computing the distance among the sample and the clustering center, a mapping *ϕ*_*k*_ is formed from *d* -dimensional space to the 2*m*_*k*_-dimensional space, and the feature subset of the label in each instance is obtained as a basis for judging whether the label exists or not, such as equation (19).

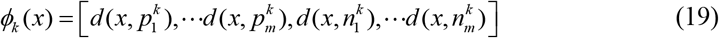

Finally, *q* classification models are constructed and recorded as {*g*_1_, *g*_2_,…, *g*_*q*_}. For each category labeled *l*_*k*_ ∊ *Y*, a binary classification training sample *B*_*k*_ containing *m* samples are created from the multi-label training sample *D* and mapping *ϕ*_*k*_, such as equation (20).

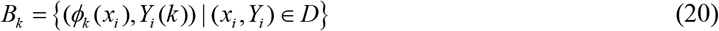

where 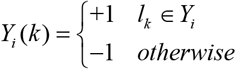. Based on the new training set, SVM is selected for classification in this study. Thus, the prediction outcomes of each category label are obtained.

### 2.5. Performance evaluation and model construction

In statistical learning, re-substitution, jackknife [50], k-fold cross-validation [51] and independent test, etc. are usually used to check out the effectiveness of the model. We use jackknife to test the performance of the model. Specifically, every time a sequence in the dataset is selected as the test sample, and others are used as the performance of the training sample training model, repeating n times until all the sequences are tested. To assess the effectiveness of the prediction model more intuitively, the overall location accuracy (OLA) and the overall actual accuracy (OAA) are used to assess the outcomes of the prediction model [52, 53]. The equations are as follows:

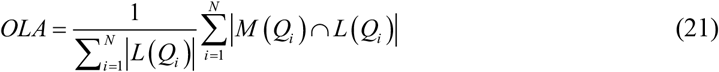

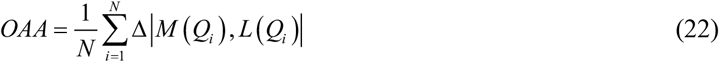

where, *L*(*Q*_*i*_) and *M*(*Q*_*i*_) represent the real label and predicted label, | · | counts the labels in the collection, ∩ is the intersection of the two collections, 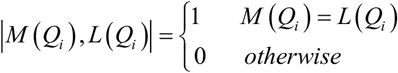.

In addition, some other performance indicators, such as Coverage (CV), Average precision (AP), Ranking Loss (RL) and Hamming Loss (HL) [54], are also used to assess the model. And indicators are defined as follows:

1. Hamming loss (HL):

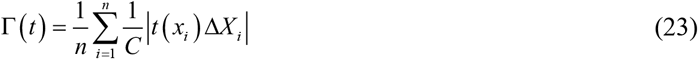
2. Coverage (CV):

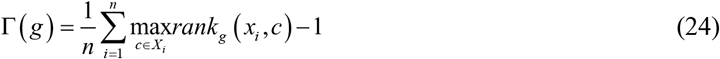
3. Ranking loss (RL):

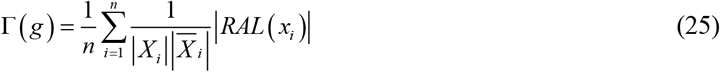

where 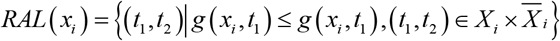.
4. Average precision (AP):

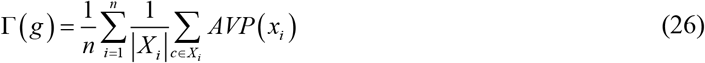

where 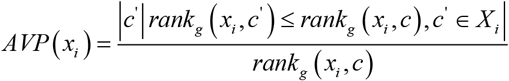.

where, Γ={(*x*_*i*_, *X*_*i*_)1 ≤ *i* ≤ *n*} represents a given multi-label training collection. For HL, CV and RL, the smaller the value, the better the performance of the model. For AP, OAA and OLA, the larger the value, the better the performance.

In our study, the protein SCL prediction method is called DMLDA-LocLIFT. The calculation process is displayed in Fig. 1. MATLAB2014a are implemented to fulfill the prediction, and the simulation environment is as follows: Windows Server 2012R2 Intel (R) Xeon (TM) CPU E5-2650 @ 2.30GHz with 32.0GB of RAM.

**Fig. 1.**
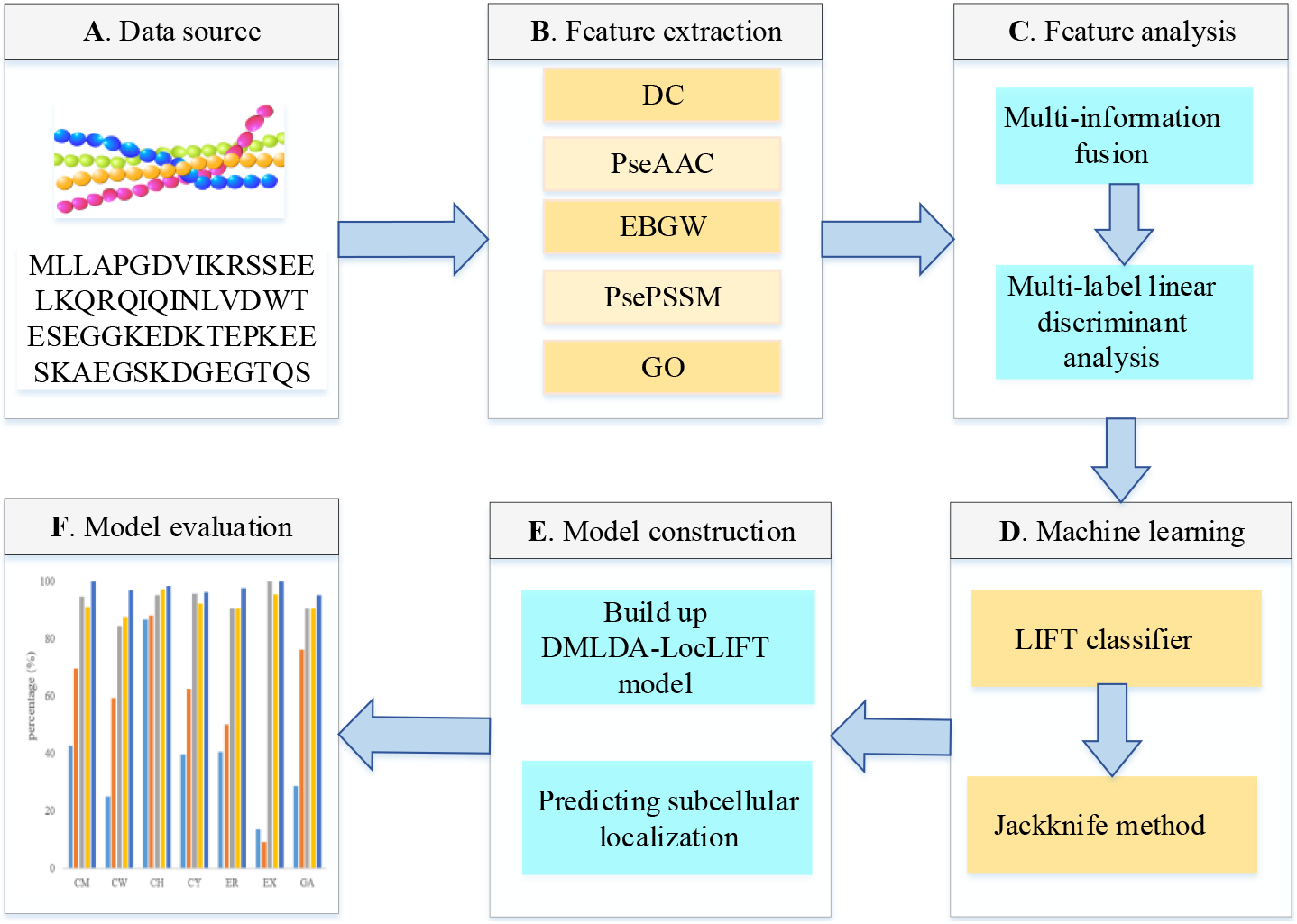
Flowchart of protein SCL used in DMLDA-LocLIFT. (A) Input Protein sequences. (B) Using five feature extraction methods to draw effective information from sequences (C) The extracted features are fused and dimensionality is reduced using the DMLDA algorithm to remove redundancy. (D) Prediction results based on the LIFT classifier and jackknife method. (E) Constructing DMLDA-LocLIFT Model and calculate the final prediction results. (F) Evaluate model performance using a test set.

The steps of DMLDA-LocLIFT are depicted as follows:

**Step 1:** Prepare protein sequences of Gram-positive bacteria and Gram-negative bacteria datasets and corresponding class labels of proteins.
**Step 2:** The PseAAC web service system is used to draw the features from protein sequences and generate 20+*λ* dimensional feature vectors; PsePSSM algorithm generates 20+20×*ξ* dimensional vectors; EBGW algorithm generates 3×*L* dimensional eigenvectors; DC algorithm generates 400-dimensional eigenvectors; For Gram-negative bacteria dataset, the GO algorithm is used to generate 1717-dimensional feature vectors; For Gram-positive bacteria dataset, the GO algorithm is used to generate 912-dimensional feature vectors. Five kinds of feature extraction information are fused and the feature matrices generated by two different bacterial datasets are *X* = (20 + *λ*) + (20+20×*ξ*) + (3×*L*) + 400 +1717 and *X* = (20 + *λ*) + (20+20 ×*ξ*) + (3×*L*) + 400 + 912 dimensions feature vectors respectively.
**Step 3:** Use DMLDA method to decrease the dimension of the fused feature in **Step 2** and LIFT is selected as the classifier. The optimal feature vectors via feature selection are determined by OAA, OLA and other evaluation indexes obtained using the jackknife test.
**Step 4:** By the jackknife test, the first-best feature subsets are input into the LIFT classifier to predict the protein SCL, and the DMLDA-LocLIFT model is determined.
**Step 5:** The OAA, OLA and other indicators of the two training sets obtained by the model are calculated to assess the prediction performance of the model.
**Step 6:** The performance of the DMLDA-LocLIFT prediction model is tested using plant dataset as independent test sets.

## 3. Results

### 3.1. Selection of optimal parameters λ, ξ and L

In this text, using the PseAAC, PsePSSM and EBGW to draw the feature from protein sequences, the built-in parameter *λ* value, *ξ* value and *L* value of the algorithm play a vital part in the model. To getting the best feature information, it is necessary to continuously debug the parameters. For the datasets of Gram-negative bacteria and Gram-positive bacteria, to seek out the best parameters for feature extraction, the *ξ* values are set to 0 to 10 in turn; the *L* values are set from 5 to 45 with an interval of 5. Since the minimum length of the selected sequence is 50 when constructing protein datasets, the *λ* values are set from 0 to 49 in turn. Tested by the jackknife method, two training datasets are predicted by the LIFT classifier, and the specific prediction results are shown in Supplementary Table S3 to S8. Fig. 2 shows the change in OAA values corresponding to different parameters selected under Gram-negative bacteria and Gram-positive bacteria datasets.

**Fig. 2.**
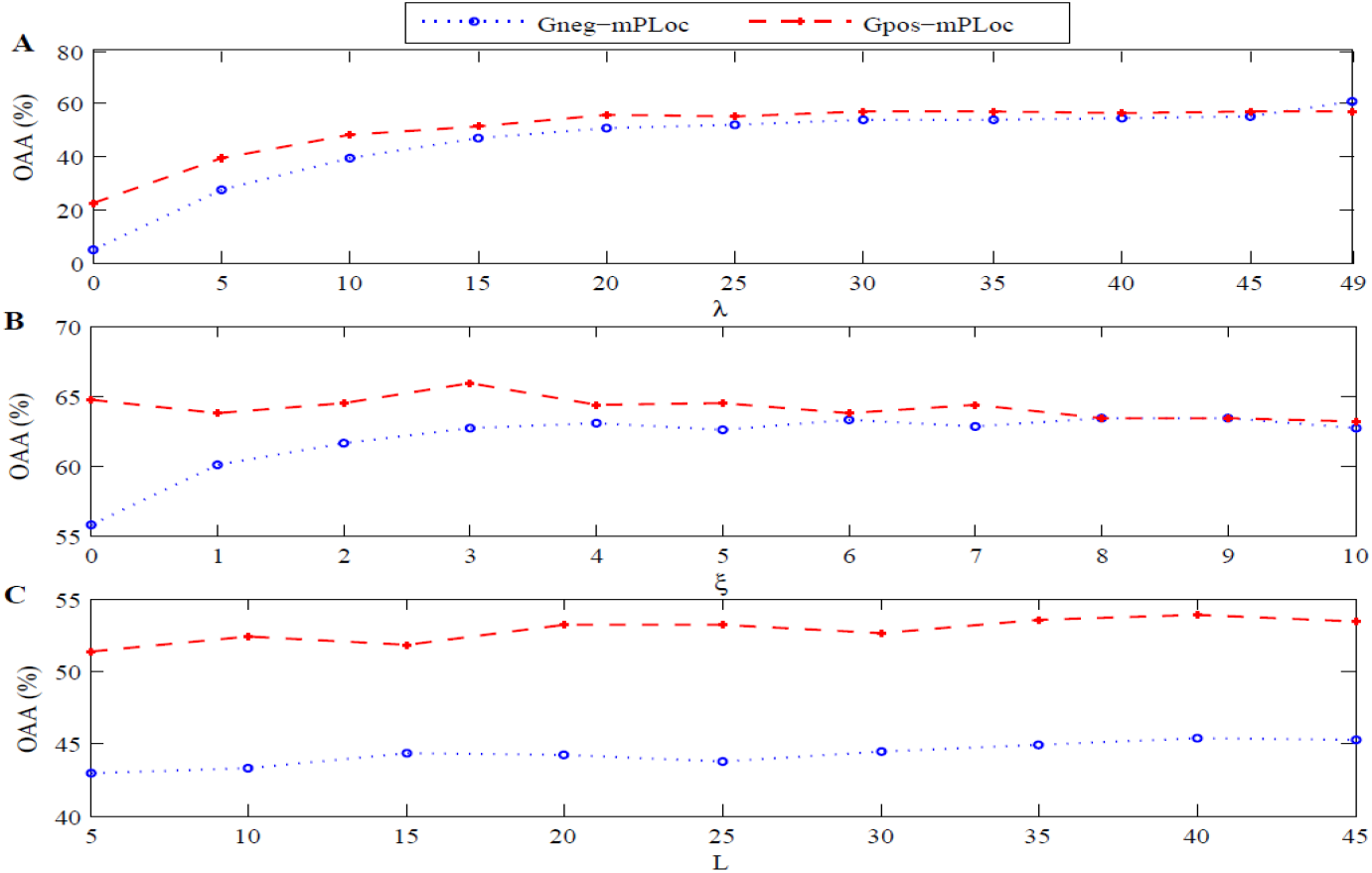
The influence of different parameters on the overall actual accuracy of Gram-negative bacteria and Gram-positive bacteria datasets. (A) Using the PseAAC algorithm to obtain the overall actual accuracy. (B) Using the PsePSSM algorithm to get the overall actual accuracy. (C) Using the EBGW algorithm to obtain the overall actual accuracy.

Fig. 2 shows that with the change of the parameters, the OAA of the two datasets also changes. When using PseAAC to feature extraction, the OAA of Gram-positive bacteria reached the highest when the *λ* value is 45. Gram-negative bacteria achieved the highest accuracy when the *λ* value is 49. To determine the parameters of three algorithms, the best value in the model is 49. Therefore, when using the PseAAC algorithm to draw features from protein sequences, every sequence can be produced 20 + *λ* = 69 dimensional feature vectors. When using the PsePSSM to extract feature, we can see when the *ξ* value of Gram-positive bacteria is 3, and the *ξ* value of Gram-negative bacteria is 9, the highest OAA value is achieved. Considering comprehensively *ξ* equals 3 is taken as the optimal parameter of the model. So, when using the PsePSSM algorithm to draw features from protein sequences, every sequence can be produced 20 + 20×*ξ* = 80 dimensional feature vectors. When using EBGW for feature extraction, Gram-negative bacteria and Gram-positive bacteria reach the highest accuracy when the *L* value is 40, so the *L* value of this paper is 40. Therefore, when the EBGW algorithm is used for drawing feature from protein sequences, every sequence can be produced 3×*L* = 120 dimensional feature vectors.

### 3.2. Effect of feature extraction algorithm on results

It is hard to obtain better prediction results and analyze biological significance only by using a single feature. More and more researchers describe protein sequences by fusing multiple features to obtain more effective feature information of protein sequences. In this paper, five single feature extraction methods will be used to obtain more feature information, but many redundant information will inevitably be produced. Therefore, the DMLDA dimensionality reduction method is used to eliminate noise and preserve important and effective feature information in protein sequences. In this paper, five separate feature extraction methods (PseAAC, PsePSSM, EBGW, DC and GO information) are used to compare the results of feature fusion based on DMLDA dimensionality reduction. PseAAC can extract frequency information, physicochemical information and sequence information of amino acids. PsePSSM can draw sequence and evolutionary information from protein sequences. EBGW draws physicochemical information from sequences. DC extracts the information in amino acid pairs of protein sequences. GO information can obtain annotation information of proteins. The LIFT classifier is selected to predict the Gram-negative bacteria and Gram-positive bacteria datasets, using the jackknife method to evaluate the datasets, and the specific prediction results about different feature extraction algorithms are obtained as shown in Supplementary Table S9. To determine the best feature extraction method, Fig. 3 shows the changes of OAA and OLA values of Gram-negative bacteria and Gram-positive bacteria with different feature extraction algorithms.

**Fig. 3.**
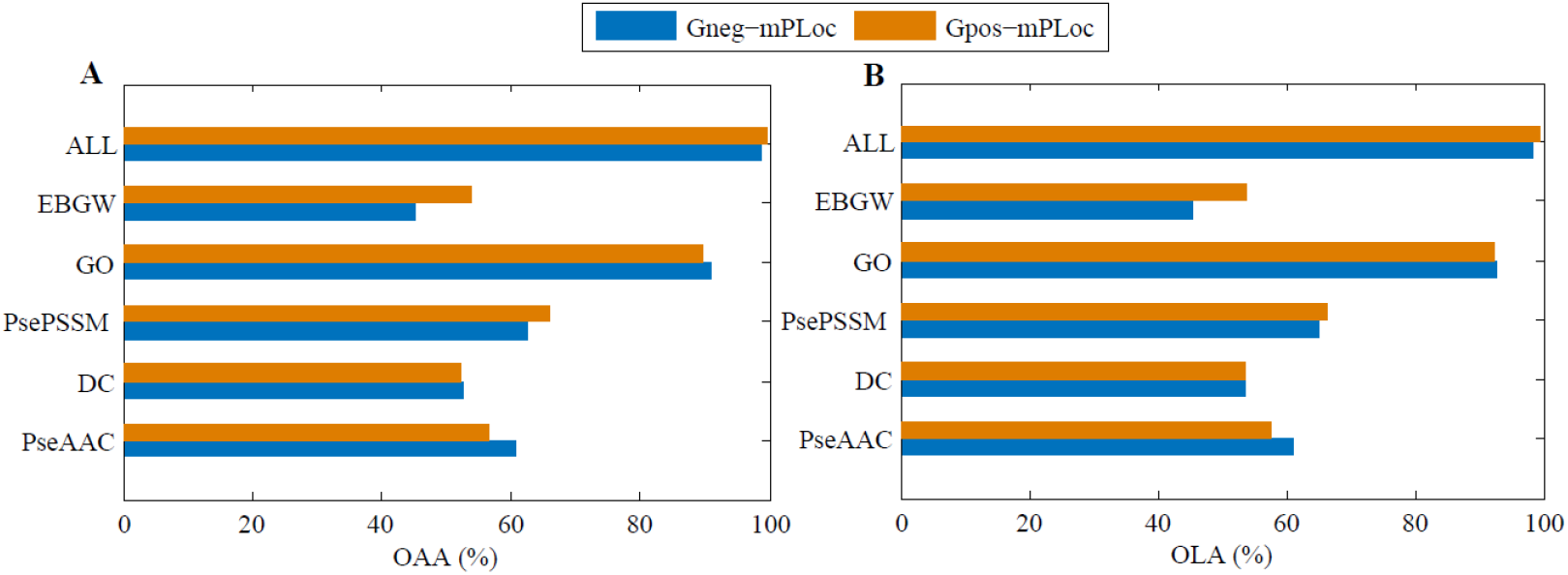
The influence about different feature extraction methods on the prediction accuracy of Gram-positive bacteria and Gram-negative bacteria. (A) The overall actual accuracy of feature extraction using different methods. (B) The overall location accuracy of feature extraction using different methods

It can be concluded from Fig. 3 that different feature extraction methods result in different prediction results on the two bacterial datasets. For single feature extraction methods, GO information has the greatest impact on multi-label protein SCL prediction. About the Gram-negative bacteria dataset, the OAA and OLA of GO information are 90.9% and 92. 8%, respectively. Among the prediction results of every protein class, GO is better than other single feature extraction methods significantly. For the feature fusion method based on DMLDA dimensionality reduction, OAA and OLA reach 98.6% and 98.6%, respectively, which are 7.7% and 5.8% higher than GO information. About the Gram-positive bacteria dataset, the prediction results are the best by using the feature fusion method based on DMLDA dimensionality reduction, and the prediction accuracy of OAA and OLA are 99.6% and 99.4%, respectively. Moreover, the prediction results of every class of proteins is significantly higher than that of other single feature extraction methods. Therefore, we choose to combine the five feature extraction algorithms, which are PseAAC, PsePSSM, EBGW, DC and GO information, and use DMLDA to remove the redundancy and noise information caused by fusion. Finally, we get better prediction results of protein SCL.

Among them, when making use of the PseAAC algorithm for feature extraction, 69-dimensional feature vectors can be produced when *λ*=49 is taken. When employing the PsePSSM algorithm for feature extraction, 80-dimensional feature vectors can be produced when *ξ*=3 is taken. When making use of the EBGW algorithm for feature extraction, 120-dimensional feature vectors can be produced when *L* = 40 is taken. A 400-dimensional feature vector is generated when protein sequences are extracted using a DC algorithm. When GO information is used for feature extraction, 1717-dimensions feature vectors are generated for Gram-negative bacteria and 912-dimensions feature vectors are generated for Gram-positive bacteria. Therefore, when using five single feature extraction algorithms to fuse, a total of 2386-dimensions feature vectors are generated for Gram-negative bacterial datasets, and 1581-dimensions feature vectors are generated for Gram-positive bacterial datasets.

### 3.3. Effect of dimensional reduction algorithm on results

For the Gram-negative bacteria and Gram-positive bacteria datasets, five feature extraction algorithms, PseAAC, PsePSSM, EBGW, dipeptide composition and GO information, are fused to draw the features from protein sequences. The feature vectors of 2386 and 1581-dimensions are obtained respectively, but it also brings a lot of redundant features. To reach the desired prediction accuracy of protein SCL, principal component analysis (PCA), multi-label informed latent semantic indexing (MLSI), Multi-label dimensionality reduction via dependency maximization (MDDM), Multi-label feature extraction algorithm via maximizing feature variance and feature-label dependence (MVMD) and Direct multi-label linear discriminant analysis (DMLDA) are used to reduce the dimension. Through LIFT multi-label classification, jackknife is applied to test the results, and the OAA value of two datasets are obtained by selecting different dimensions under different dimensionality reduction methods, as shown in Supplementary Table S10. To more intuitively analyze the predicted performance of Gram-negative bacteria and Gram-positive bacteria under different dimensionality reduction methods, Fig. 4 and Fig. 5 show the comparison results of OAA, HL, CV and AP in different dimensions.

**Fig. 4.**
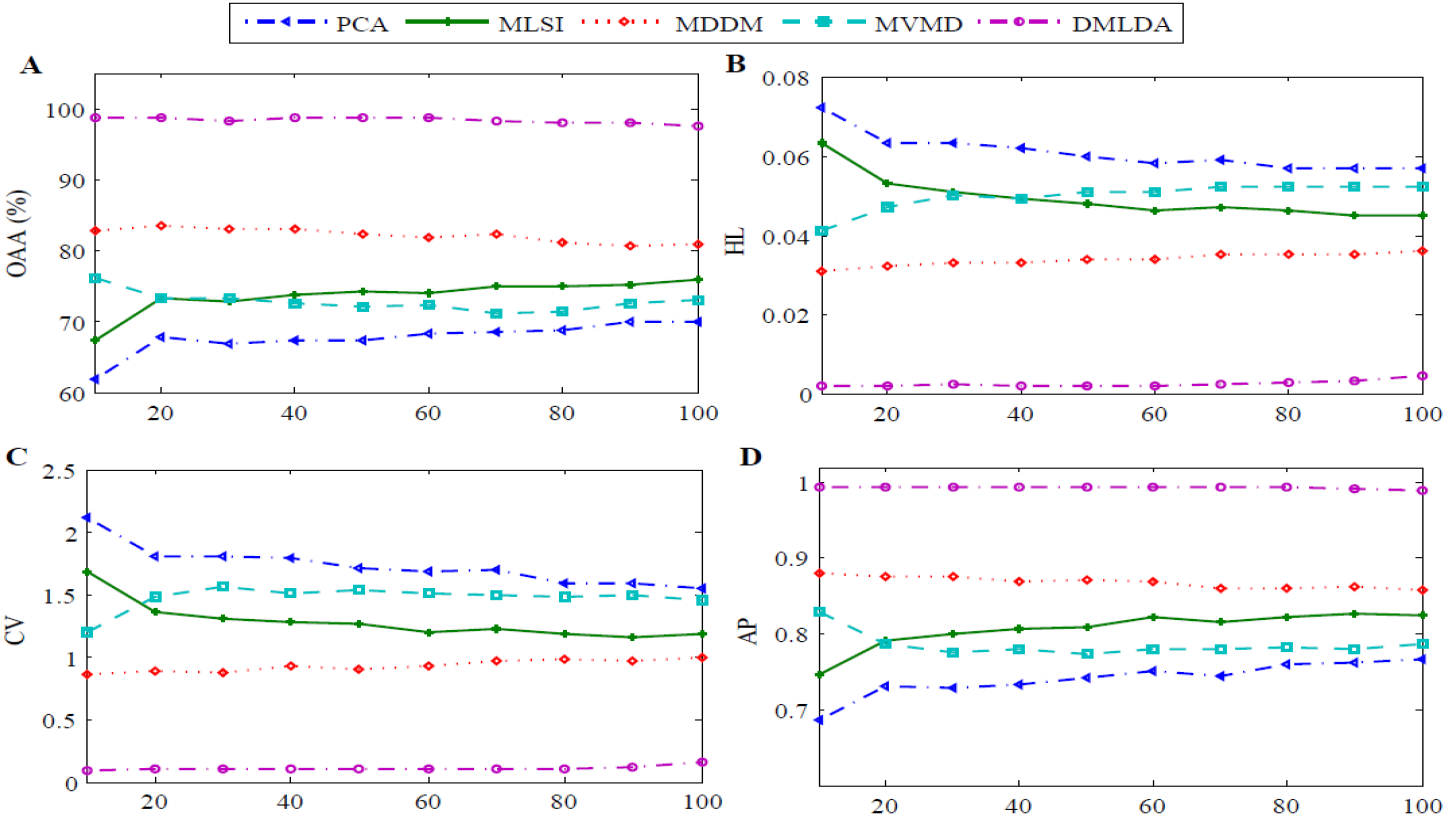
The effect of selecting different dimensions under different dimensionality reduction methods on the predictive results of Gram-negative bacteria. (A) The overall actual accuracy under different methods and dimensions. (B) The hamming loss under different methods and dimensions. (C) The coverage under different methods and dimensions. (D) The average precision under different methods and dimensions

**Fig. 5.**
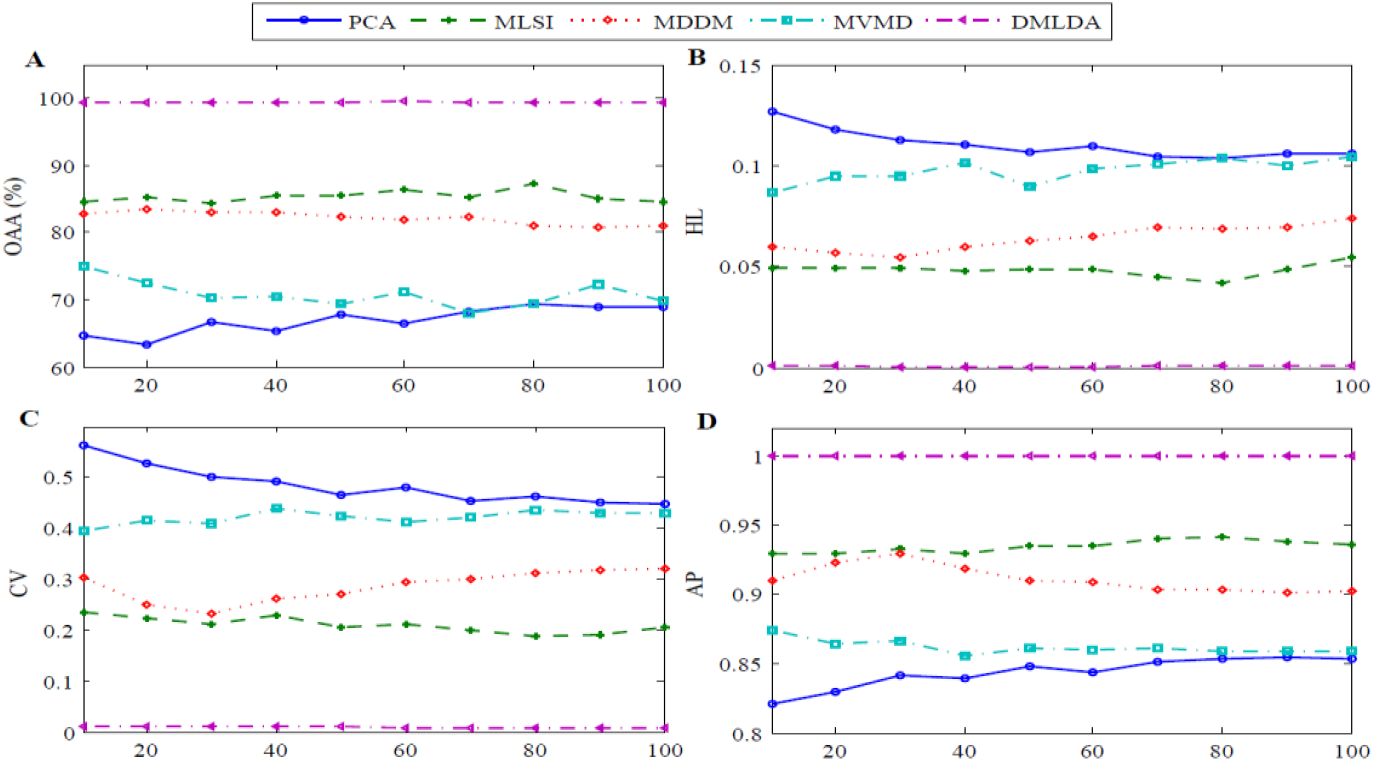
The effect of selecting different dimensions under different dimensionality reduction methods on the predictive results of Gram-positive bacteria. (A) The overall actual accuracy under different methods and dimensions. (B) The hamming loss under different methods and dimensions. (C) The coverage under different methods and dimensions. (D) The average precision under different methods and dimensions.

In Fig. 4 and Fig. 5, selecting different dimension reduction methods and dimensions has great impact on the OAA of protein subcellular prediction and other indicators. By comparing the five dimensionality reduction methods of PCA, MLSI, MDDM, MVMD and DMLDA, for the Gram-negative dataset, when DMLDA is used to reduce dimension and 10 features are selected as the feature subset, the OAA reaches 98.7%, which is 36.9%, 31.4%, 15.9%, and 22.6% higher than that of PCA, MLSI, MDDM and MVMD, respectively, which is obviously superior to other dimension reduction methods when they take different dimensions. Similarly, for the Gram-positive dataset, when DMLDA is used to reduce dimension and 60 features are selected as the feature subset, the OAA reaches 99.6%, which is 33.3%, 13.3%, 17.7%, and 28.5% higher than that of PCA, MLSI, MDDM and MVMD, respectively, which is obviously superior to other dimension reduction methods when they take different dimensions. For two bacterial datasets, the DMLDA method achieves the best performance under Hamming loss, coverage, and other indicators. By comparing the results, we use the DMLDA dimensionality reduction method and the dimension is 60-dimensions. The overall accuracy of the model is the highest.

### 3.4. Selection of classification algorithms

To construct an effective prediction model for multi-label protein SCL, the appropriate classifier algorithm has a big influence on the performance about the model. In this paper, combining five feature extraction algorithms, using DMLDA algorithm to select the first-best feature subset, and four classification algorithms, ML-KNN, ML-LOC, INSDIF and LIFT, are selected to predict the accuracy rate of Gram-negative bacteria and Gram-positive bacteria datasets. Among them, the ML-KNN classification algorithm chooses *k* = 1. The eigenvectors after dimensionality reduction by DMLDA are input into ML-KNN, ML-LOC, INSDIF, and LIFT classifiers respectively. Tested by the jackknife method, the specific outcomes of two datasets are shown in Supplementary Table S11 and S12. respectively. Fig. 6 shows the changes in six performance indexes of Gram-negative bacteria and Gram-positive bacteria with different classifier algorithms.

**Fig. 6.**
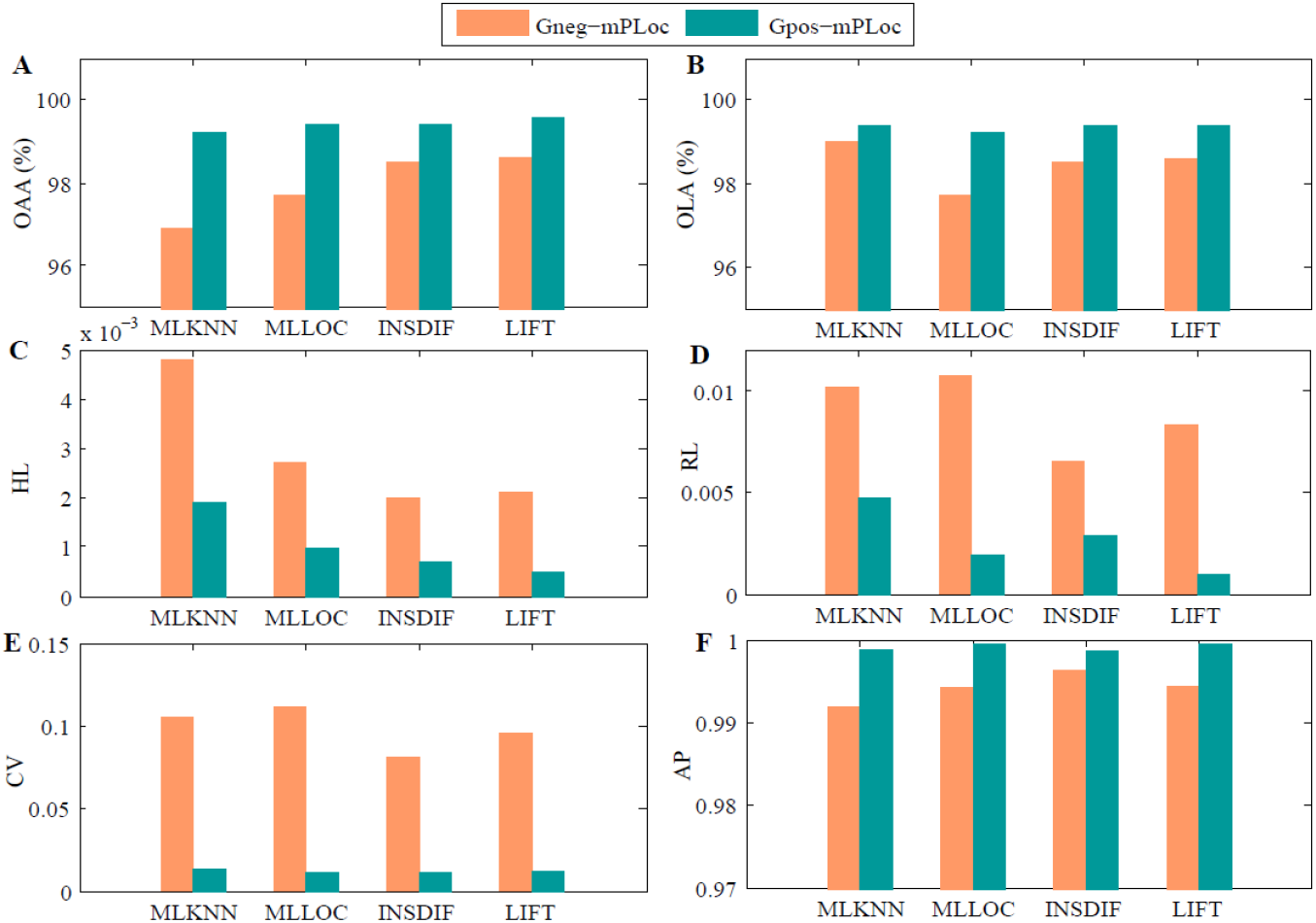
The effect of different classifiers on the prediction results of the datasets of Gram-negative bacteria and Gram-positive bacteria. (A) Using different classifiers to obtain the overall actual accuracy. (B) Using different classifiers to obtain overall location accuracy. (C) Using different classifiers to obtain hamming loss. (D) Using different classifiers to obtain the ranking loss. (E) Using different classifiers to obtain coverage. (F) Using different classifiers to obtain average precision

In Fig. 6, for the Gram-negative bacteria dataset, using the LIFT algorithm as the classifier of the model, the OLA and OAA values of the model are 98.6%, which are 0.9% higher than those of ML-LOC. Although the OLA value of ML-KNN is 0.4% higher than LIFT and OLA value of INSDIF is 0.2% higher than LIFT, the OAA value of LIFT is 1.7% and 0.3% higher than ML-KNN and INSDIF, respectively. Therefore, the average results of evaluation indexes using the LIFT classifier is better than other classifiers. For the Gram-positive bacteria dataset, when we use the LIFT algorithm as the classifier of the model, the OLA and OAA of the model reach 99.4% and 99.6% respectively, which are 0.4% higher than ML-KNN’s OAA value and 0.2% higher than INSDIF’s OAA value. The outcomes are significantly superior to other classification algorithms. By comparing the results of Hamming loss and average accuracy, the performance of the LIFT is superior to other classifiers for datasets of Gram-negative bacteria and Gram-positive bacteria.

Through the analysis of OLA, OAA and other performance indicators of two datasets on four classifiers, a prediction model with high stability is selected. For the two training sets in this paper, the ML-KNN algorithm needs a lot of space to store instances, which has high complexity and unstable prediction performance. Although ML-LOC and INSDIF have considerable prediction results, they are running slowly and still have rising space. Because LIFT changes the multi-label question into the single-label question, and label information in the dataset is fully utilized. Through the clustering analysis of each label, the characteristics of the label are constructed, and the instances are trained and tested by the clustering results. Considering comprehensively, we choose the LIFT classification algorithm as the model classification algorithm.

### 3.5 Comparison with other methods

At present, researchers have put forward many methods to predict protein SCL of Gram-negative bacteria and Gram-positive bacteria datasets. To prove the validity of the model proposed in this paper, we contrast its prediction results to the identical dataset based on the jackknife test. Firstly, PSeAAC, EBGW, PSePSSM, GO and DC are used to extract and fuse the features, and DMLDA is applied to select the optimal feature subset, and then using LIFT classifier to obtain the prediction results. The comparative results of Gram-negative bacteria and Gram-positive bacteria datasets are shown in Table 2 and Table 3.

**Table 2.**
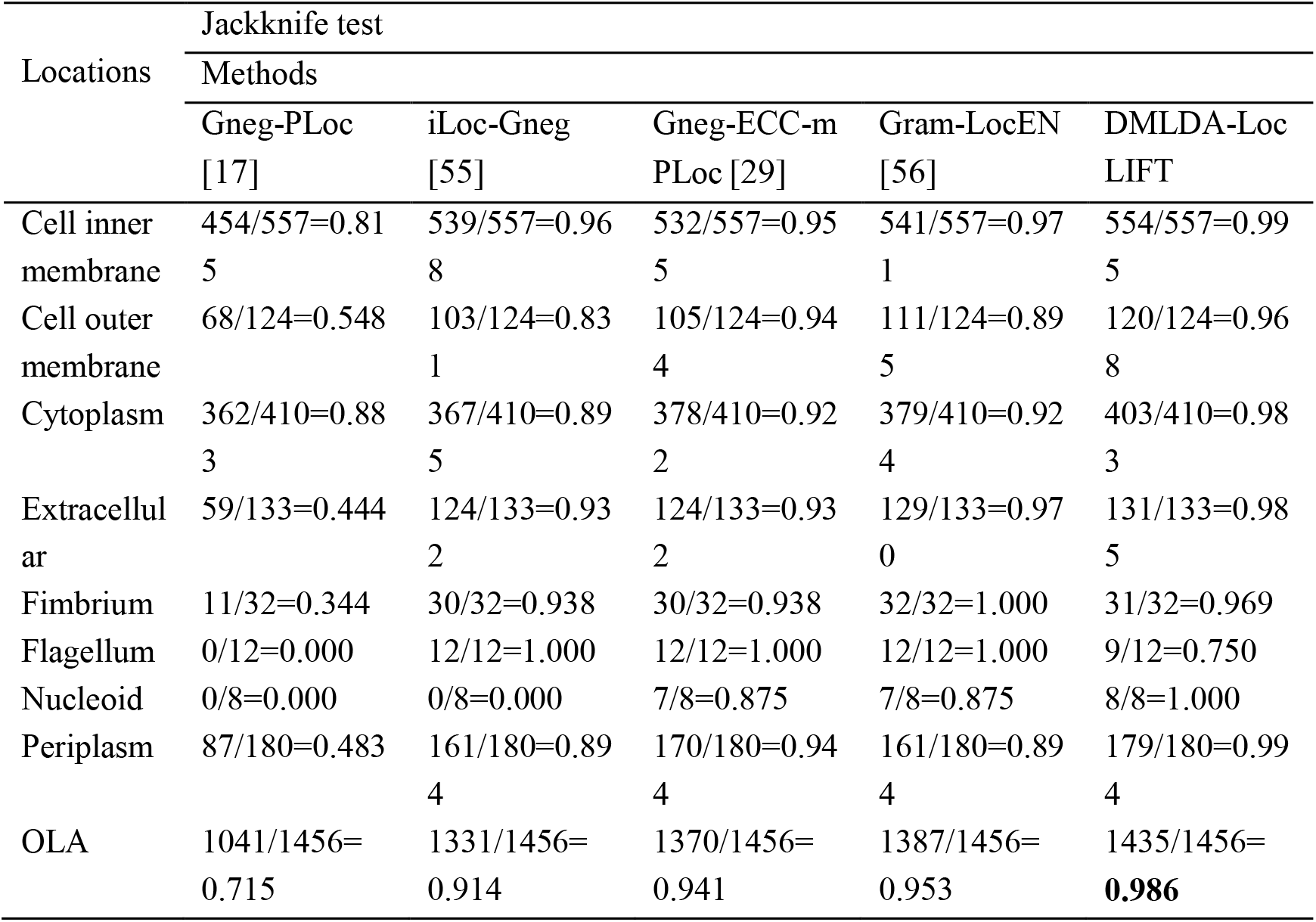
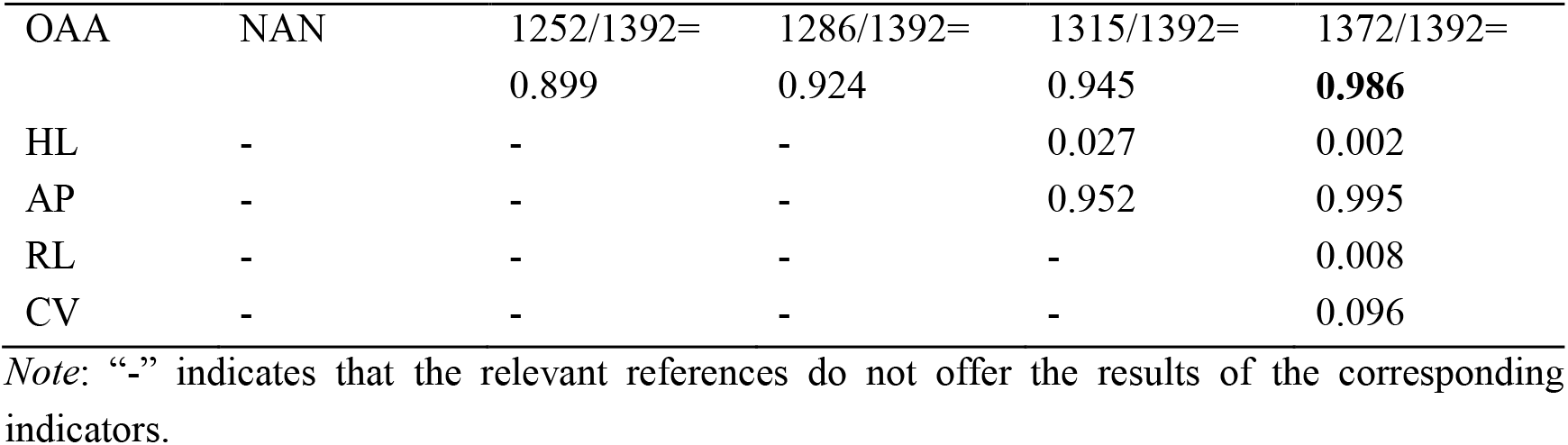
Comparison of the results of using different prediction methods on Gram-negative bacteria.

**Table 3.**
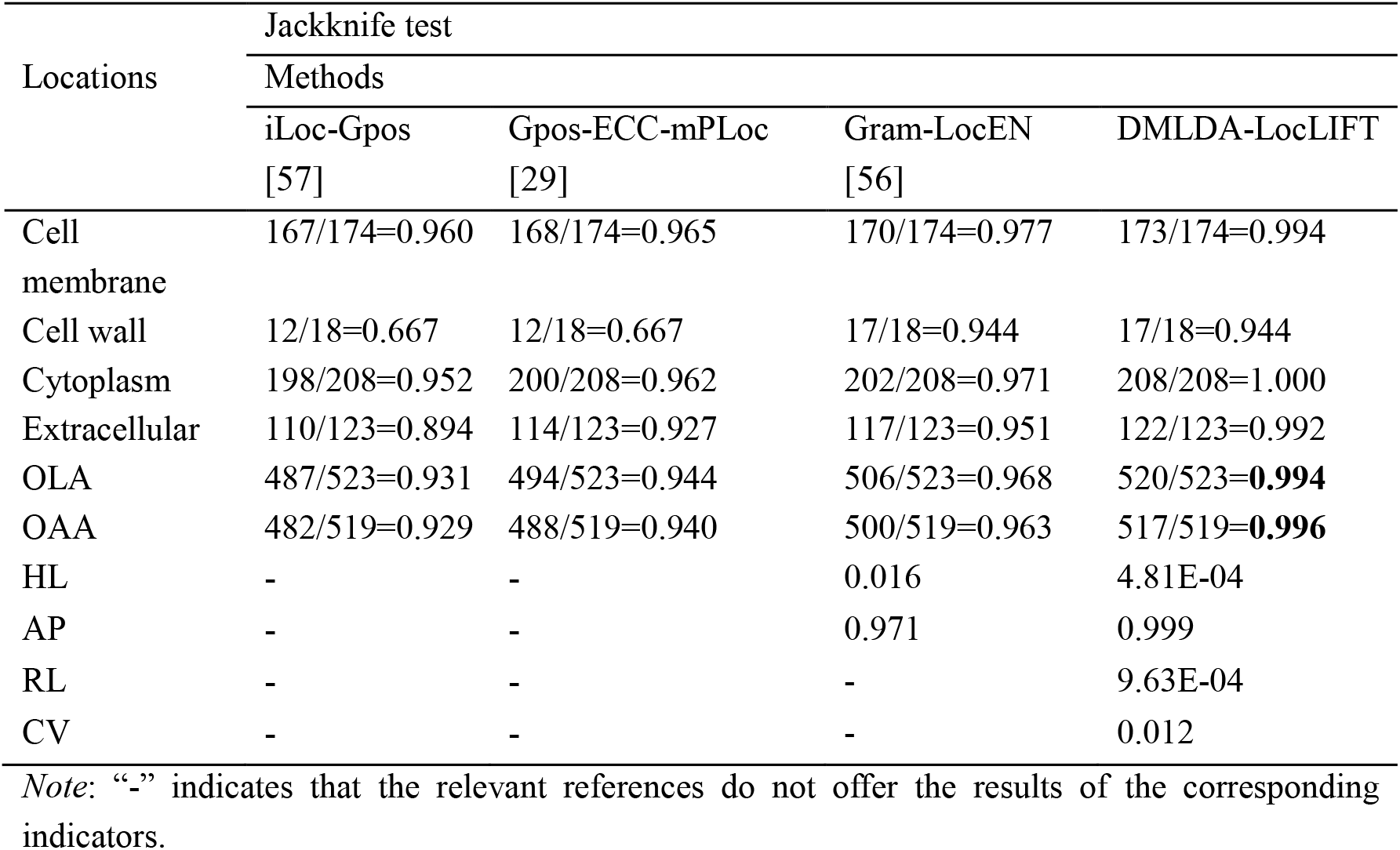
Comparison of the results of using different prediction methods on Gram-positive bacteria.

| Locations | Jackknife test |  |  |  |
| --- | --- | --- | --- | --- |
|  | Methods |  |  |  |
|  | iLoc-Gpos<br>[57] | Gpos-ECC-mPLoc<br>[29] | Gram-LocEN<br>[56] | DMLDA-LocLIFT |
| Cell membrane | 167/174=0.960 | 168/174=0.965 | 170/174=0.977 | 173/174=0.994 |
| Cell wall | 12/18=0.667 | 12/18=0.667 | 17/18=0.944 | 17/18=0.944 |
| Cytoplasm | 198/208=0.952 | 200/208=0.962 | 202/208=0.971 | 208/208=1.000 |
| Extracellular | 110/123=0.894 | 114/123=0.927 | 117/123=0.951 | 122/123=0.992 |
| OLA | 487/523=0.931 | 494/523=0.944 | 506/523=0.968 | 520/523=**0.994** |
| OAA | 482/519=0.929 | 488/519=0.940 | 500/519=0.963 | 517/519=**0.996** |
| HL | - | - | 0.016 | 4.81E-04 |
| AP | - | - | 0.971 | 0.999 |
| RL | - | - | - | 9.63E-04 |
| CV | - | - | - | 0.012 |

*Note:* “-” indicates that the relevant references do not offer the results of the corresponding indicators.

In Table 2 that for the Gram-negative bacteria dataset, the OLA of the proposed model DMLDA-LocLIFT is 98.6%, which is 3.3%-27.1% higher than other prediction algorithms. The OAA is 98.6%, which is 4.1% −8.7% higher than other prediction algorithms. For Nucleoid protein, the prediction accuracy of the DMLDA-LocLIFT algorithm is 100%, while that of Gneg-PLoc algorithm and the iLoc-Gneg algorithm is 0%. The prediction accuracy of Gneg-ECC-mPLoc and Gram-LocEN algorithms are 87.5%. This shows that our method definitely increases the accuracy of Nucleoid protein. For Periplasm protein, the accuracy of this algorithm is 99.4%, which is 5%-51.1% better than other prediction methods. Although the prediction performance of the DMLDA-LocLIFT algorithm is slightly worse for Fimbrium and Flagellum proteins, it has been significantly improved at other locations. Considering comprehensively, in this paper, DMLDA-LocLIFT has a good performance on the dataset of Gram-negative bacteria.

In Table 3 that for the Gram-positive bacteria dataset, the OLA of the proposed model DMLDA-LocLIFT is 99.4%, which is 2.6%-6.3% higher than other methods. The OAA is 99.6%, which is 3.3%-6.7% higher than other prediction algorithms. Moreover, for each site, the accuracy of the DMLDA-LocLIFT model is superior to other prediction methods. By comparison, it is found that the DMLDA-LocLIFT model proposed in this paper has achieved satisfactory results.

To verify the predictive capability about the model, we use plant dataset as the independent test set, and its results are compared with other methods in Table 4, and the comparison chart between the methods is shown in Supplementary Fig. S1. For the plant dataset, the OLA of the proposed model DMLDA-LocLIFT is 97.5%, which is 1.3%-33.8% higher than other prediction algorithms. The OAA is 97.9%, which is 4.3%-29.8% better than other methods. For Cell membrane protein, Extracellular protein and Peroxisome protein, the accuracy of the DMLDA-LocLIFT is 100%, which is significantly higher than that of other algorithms. For Plastid protein, although the predictive effect of the DMLDA-LocLIFT algorithm is slightly worse than that of the HybridGO-Loc and mGOASVM algorithm, the predictive results of this model are better than those of other prediction algorithms at other sites. Especially in the prediction of Cytoplasm protein, Endoplasmic reticulum protein and Vacuole protein, the prediction accuracy of the DMLDA-LocLIFT algorithm is 3.9%, 7.1%, and 5.8% higher than that of HybridGO-Loc algorithm, respectively. Considering comprehensively, the proposed method has good prediction performance on plant dataset.

**Table 4.**
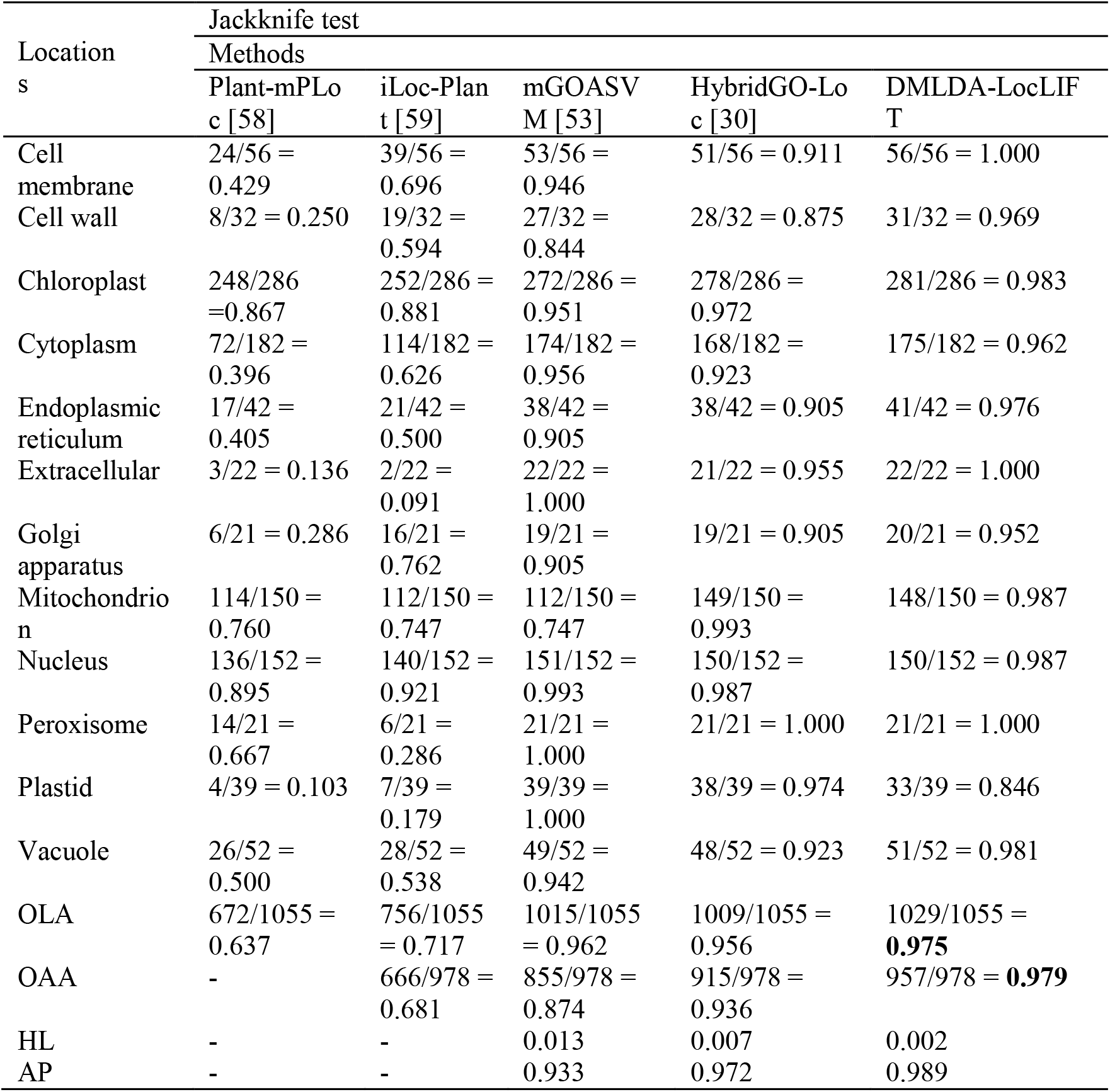
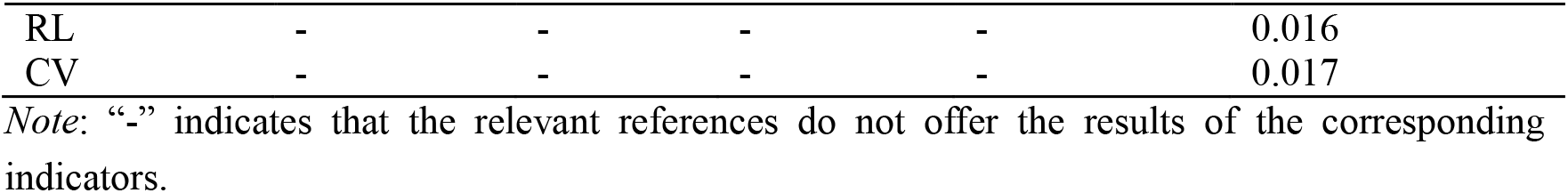
Comparison of different prediction methods for protein subcellular localization on plant dataset.

| Location<br>s | Jackknife test |  |  |  |  |
| --- | --- | --- | --- | --- | --- |
|  | Methods |  |  |  |  |
|  | Plant-mPLOC [58] | iLoc-Plant [59] | mGOASVM [53] | HybridGO-Loc [30] | DMLDA-LocLIFT |
| Cell membrane | 24/56 = 0.429 | 39/56 = 0.696 | 53/56 = 0.946 | 51/56 = 0.911 | 56/56 = 1.000 |
| Cell wall | 8/32 = 0.250 | 19/32 = 0.594 | 27/32 = 0.844 | 28/32 = 0.875 | 31/32 = 0.969 |
| Chloroplast | 248/286 = 0.867 | 252/286 = 0.881 | 272/286 = 0.951 | 278/286 = 0.972 | 281/286 = 0.983 |
| Cytoplasm | 72/182 = 0.396 | 114/182 = 0.626 | 174/182 = 0.956 | 168/182 = 0.923 | 175/182 = 0.962 |
| Endoplasmic reticulum | 17/42 = 0.405 | 21/42 = 0.500 | 38/42 = 0.905 | 38/42 = 0.905 | 41/42 = 0.976 |
| Extracellular | 3/22 = 0.136 | 2/22 = 0.091 | 22/22 = 1.000 | 21/22 = 0.955 | 22/22 = 1.000 |
| Golgi apparatus | 6/21 = 0.286 | 16/21 = 0.762 | 19/21 = 0.905 | 19/21 = 0.905 | 20/21 = 0.952 |
| Mitochondrion | 114/150 = 0.760 | 112/150 = 0.747 | 112/150 = 0.747 | 149/150 = 0.993 | 148/150 = 0.987 |
| Nucleus | 136/152 = 0.895 | 140/152 = 0.921 | 151/152 = 0.993 | 150/152 = 0.987 | 150/152 = 0.987 |
| Peroxisome | 14/21 = 0.667 | 6/21 = 0.286 | 21/21 = 1.000 | 21/21 = 1.000 | 21/21 = 1.000 |
| Plastid | 4/39 = 0.103 | 7/39 = 0.179 | 39/39 = 1.000 | 38/39 = 0.974 | 33/39 = 0.846 |
| Vacuole | 26/52 = 0.500 | 28/52 = 0.538 | 49/52 = 0.942 | 48/52 = 0.923 | 51/52 = 0.981 |
| OLA | 672/1055 = 0.637 | 756/1055 = 0.717 | 1015/1055 = 0.962 | 1009/1055 = 0.956 | 1029/1055 = **0.975** |
| OAA | - | 666/978 = 0.681 | 855/978 = 0.874 | 915/978 = 0.936 | 957/978 = **0.979** |
| HL | - | - | 0.013 | 0.007 | 0.002 |
| AP | - | - | 0.933 | 0.972 | 0.989 |
| RL | - | - | - | - | 0.016 |
| CV | - | - | - | - | 0.017 |

*Note:* “-” indicates that the relevant references do not offer the results of the corresponding indicators.

## 4. Discussion

In this study, we put forward a new multi-label protein prediction algorithm called DMLDA-LocLIFT. A total of five feature extraction algorithms are adopted, namely PseAAC, PsePSSM, EBGW, DC and GO. Using DMLDA algorithm to select the best-first feature subset in three datasets, the dimension is 60 after comparison. And the LIFT classifier is used to predict the position of the multi-label proteins. Through the jackknife test, the OLA of Gram-negative bacteria, Gram-positive bacteria, and plant datasets are 98.6%, 99.4%, 97.5%, which are 3.3%-27.1%, 2.6%-6.3%, 1.3%-33.8% superior to other advanced methods respectively. And OAA of three datasets are 98.6%, 99.6% and 97.9%, which are 4.1%-8.7%, 3.3%-6.7%, 4.3%-29.8% better than other advanced methods respectively.

The DMLDA-LocLIFT model can effectively predict SCL of multi-label proteins, and its precision obviously superior to other advanced methods. For the following reasons:

1. The physical and chemical information, evolutionary information, sequence information and annotation information in the fusion protein sequence are extracted by feature extraction. GO information has the highest prediction accuracy and plays a vital part in the multi-label prediction model.
2. DMLDA dimensionality reduction method can eliminate noise information in high-dimensional data using label correlation, and effectively retain the important feature vectors for recognizing protein subcellular locations.
3. By introducing the idea of clustering, LIFT converts the multi-label classification issue to multiple binary classification questions, which greatly improves the prediction accuracy of multi-locus learning, can effectively predict the subcellular location of multi-label proteins and has good model generalization ability.

To explore the SCL of multi-label proteins can help people understand the function of these proteins more accurately, and provide theoretical basis for the pathogenesis of diseases. And understanding the location of proteins can furnish new opinions with drug design and drug target identification, so as to defeat the disease.

## 5. Conclusion

Studies have found that multi-label proteins carry more cellular metabolic functions, and normal life activities within cells are more dependent on them. Through the study of multi-label protein SCL can help people find out the pathogenesis of more diseases and reveal the special functions of multi-label proteins in drug compounds. Facing with the increasing data of multi-label protein SCL, traditional research methods are expensive and time-consuming, how to predict multi-label protein subcellular using machine learning method is becoming more and more important. A new method of protein SCL prediction based on multi-label learning called DMLDA-LocLIFT is proposed in this paper. Firstly, use PseAAC, PsePSSM, EBGW, DC and GO algorithms to draw the valid information from sequences and fuse them. Then the feature fusion information is processed by DMLDA dimensionality reduction. Finally, the best-first feature subsets are input into the LIFT classifier to predict the location on Gram-negative bacteria, Gram-positive bacteria and plant datasets. Through the jackknife test, the OLA of three datasets are 98.6%, 99.4%, 97.5%, and OAA are 98.6%, 99.6% and 97.9%, respectively. By comparison, we propose the DMLDA-LocLIFT method which obviously increase the prediction precision of multi-label protein SCL, but there is still aspect for improvement in the construction of the model. Deep learning can use more data for further development, compared with traditional machine learning methods, deep learning no longer performs artificial feature extraction on data, and has the advantage of strong adaptability, easy to transform, with good model generalization ability. Predicting multi-label SCL using deep learning is our next research direction.

## Acknowledgments

This work was supported by the National Nature Science Foundation of China (No. 61863010), the Key Research and Development Program of Shandong Province of China (No. 2019GGX101001), and the Natural Science Foundation of Shandong Province of China (Nos. ZR2018MC007, ZR2019MEE066).

