## Supplementary Tables, Supplementary Figures for "DMLDA-LocLIFT: Identification of multi-label protein subcellular localization using DMLDA dimensionality reduction and LIFT classifier"

Table of Contents

### 1. Supplementary Tables

**Table S1** Breakdown of the gram-negative bacterial benchmark dataset.

**Table S2** Breakdown of the gram-positive bacterial benchmark dataset.

**Table S3** Select different parameter $\lambda$ to obtain prediction results of the Gram-negative dataset.

**Table S4** Select different parameter $\lambda$ to obtain prediction results of the Gram-positive dataset.

**Table S5** Select different parameter $\xi$ to obtain prediction results of the Gram-negative dataset.

**Table S6** Select different parameter $\xi$ to obtain prediction results of the Gram-positive dataset.

**Table S7** Select different parameter $L$ to obtain prediction results of the Gram-negative dataset.

**Table S8** Select different parameter $L$ to obtain prediction results of the Gram-positive dataset.

**Table S9** Predicting the results of different feature extraction methods for Gram-positive and Gram-negative bacteria data sets.

**Table S10** Selecting different dimensionality reduction methods and different dimensions to obtain the overall actual accuracy results of subcellular localization.

**Table S11** Prediction results of Gram-negative dataset protein subcellular localization in different classification algorithms.

**Table S12** Prediction results of Gram-positive dataset protein subcellular localization in different classification algorithms.

### 2. Supplementary Figures

**Fig. S1.** Comparison of different prediction methods for protein subcellular localization on plant dataset.

### 1. Supplementary Tables

**Table S1**

Breakdown of the gram-negative bacterial benchmark dataset. The homology of protein sequences in the dataset is limited to 25%.

| Subset | Subcellular location name | Number of proteins |
|---|---|---|
| S1 | Cell inner membrane | 557 |
| S2 | Cell outer membrane | 124 |
| S3 | Cytoplasm | 410 |
| S4 | Extracellular | 133 |
| S5 | Fimbrium | 32 |
| S6 | Flagellum | 12 |
| S7 | Nucleoid | 8 |
| S8 | Periplasm | 180 |
| Total number of locative proteins | | 1456 |
| Total number of different proteins | | 1392 |

**Table S2**

Breakdown of the gram-positive bacterial benchmark dataset. The homology of protein sequences in the dataset is limited to 25%.

| Subset | Subcellular location name | Number of proteins |
|---|---|---|
| S1 | Cell membrane | 174 |
| S2 | Cell wall | 18 |
| S3 | Cytoplasm | 208 |
| S4 | Extracellular | 123 |
| Total number of locative proteins | | 523 |
| Total number of different proteins | | 519 |

**Table S3**

Select different parameter $\lambda$ to obtain prediction results of the Gram-negative dataset.

| Locations | Jackknife method (%) | | | | | | | | | | |
|---|---|---|---|---|---|---|---|---|---|---|---|
| | $\lambda$ | | | | | | | | | | |
| | 0 | 5 | 10 | 15 | 20 | 25 | 30 | 35 | 40 | 45 | 49 |
| Inner | 0.2 | 35.4 | 53.5 | 66.1 | 71.5 | 72.2 | 73.4 | 74.5 | 75.0 | 75.8 | 75.9 |
| Outer | 16.1 | 19.4 | 22.6 | 25.8 | 25.0 | 28.2 | 32.3 | 27.4 | 29.8 | 31.5 | 33.1 |
| Cytoplasm | 4.6 | 35.1 | 48.3 | 54.6 | 59.8 | 62.4 | 63.9 | 65.4 | 65.9 | 67.1 | 71.5 |
| Extracellular | 9.8 | 11.3 | 14.3 | 12.0 | 13.5 | 12.8 | 14.3 | 15.0 | 15.0 | 15.0 | 16.5 |
| Fimbrium | 0.0 | 0.0 | 3.1 | 3.1 | 3.1 | 0.0 | 3.1 | 0.0 | 0.0 | 0.0 | 0.0 |

| | | | | | | | | | | | |
|---|---|---|---|---|---|---|---|---|---|---|---|
| Flagellum | 25.0 | 33.3 | 25.0 | 33.3 | 25.0 | 33.3 | 25.0 | 41.7 | 8.3 | 33.3 | 8.3 |
| Nucleoid | 0.0 | 0.0 | 0.0 | 0.0 | 0.0 | 0.0 | 0.0 | 0.0 | 0.0 | 0.0 | 0.0 |
| Periplasm | 15.0 | 19.4 | 22.2 | 25.6 | 27.2 | 27.8 | 29.4 | 30.0 | 30.6 | 31.7 | 61.1 |
| OLA | 5.7 | 28.8 | 40.3 | 47.5 | 51.2 | 52.5 | 54.1 | 54.7 | 55.0 | 56.1 | 61.1 |
| OAA | 4.9 | 27.7 | 39.6 | 46.9 | 50.4 | 51.7 | 53.8 | 54.1 | 54.6 | 55.4 | 60.6 |

**Table S4**

Select different parameter $\lambda$ to obtain prediction results of the Gram-positive dataset.

| Locations | Jackknife method (%) | | | | | | | | | | |
|---|---|---|---|---|---|---|---|---|---|---|---|
| | $\lambda$ | | | | | | | | | | |
| | 0 | 5 | 10 | 15 | 20 | 25 | 30 | 35 | 40 | 45 | 49 |
| Inner | 24.7 | 42.0 | 49.4 | 49.4 | 50.0 | 50.6 | 51.1 | 51.7 | 51.7 | 53.4 | 50.0 |
| Outer | 5.6 | 5.6 | 0.0 | 0.0 | 11.1 | 5.6 | 5.6 | 16.7 | 0.0 | 0.0 | 5.6 |
| Cytoplasm | 18.3 | 40.9 | 55.3 | 62.0 | 67.8 | 68.8 | 71.2 | 73.1 | 71.6 | 73.1 | 73.6 |
| Extracellular | 29.3 | 41.5 | 44.7 | 46.3 | 51.2 | 48.8 | 50.4 | 46.3 | 48.8 | 48.8 | 48.8 |
| OLA | 22.6 | 40.2 | 48.9 | 52.0 | 56.0 | 55.8 | 57.4 | 57.7 | 57.2 | 58.3 | 57.6 |
| OAA | 22.2 | 39.3 | 48.4 | 51.3 | 55.5 | 55.1 | 56.8 | 56.8 | 56.1 | 57.0 | 56.8 |

**Table S5**

Select different parameter $\xi$ to obtain prediction results of the Gram-negative dataset.

| Locations | Jackknife method (%) | | | | | | | | | | |
|---|---|---|---|---|---|---|---|---|---|---|---|
| | $\xi$ | | | | | | | | | | |
| | 0 | 1 | 2 | 3 | 4 | 5 | 6 | 7 | 8 | 9 | 10 |
| Inner | 74.3 | 75.0 | 75.6 | 75.6 | 75.4 | 75.6 | 76.1 | 75.9 | 75.8 | 75.6 | 75.6 |
| Outer | 15.3 | 31.5 | 37.9 | 37.1 | 37.1 | 37.1 | 35.5 | 34.7 | 36.3 | 36.3 | 36.3 |
| Cytoplasm | 67.8 | 72.4 | 73.9 | 76.3 | 77.1 | 78.0 | 79.8 | 79.5 | 80.7 | 80.5 | 80.7 |
| Extracellular | 18.8 | 30.8 | 39.1 | 41.4 | 41.4 | 41.4 | 41.4 | 39.8 | 36.8 | 42.9 | 40.6 |
| Fimbrium | 12.5 | 18.8 | 28.1 | 31.3 | 28.1 | 34.4 | 21.9 | 15.6 | 21.9 | 25.0 | 9.4 |
| Flagellum | 58.3 | 75.0 | 75.0 | 91.7 | 91.7 | 91.7 | 91.7 | 91.7 | 83.3 | 91.7 | 83.3 |
| Nucleoid | 0.0 | 0.0 | 0.0 | 0.0 | 0.0 | 0.0 | 0.0 | 0.0 | 0.0 | 0.0 | 0.0 |
| Periplasm | 43.3 | 50.0 | 48.9 | 50.0 | 50.6 | 47.8 | 47.2 | 47.2 | 48.9 | 47.8 | 47.2 |
| OLA | 56.7 | 61.8 | 63.8 | 65.0 | 65.1 | 65.2 | 65.5 | 65.0 | 65.4 | 65.8 | 65.2 |
| OAA | 55.8 | 60.1 | 61.6 | 62.7 | 63.0 | 62.6 | 63.3 | 62.8 | 63.4 | 63.4 | 62.7 |

**Table S6**

Select different parameter $\xi$ to obtain prediction results of the Gram-positive dataset.

| Locations | Jackknife method (%) | | | | | | | | | | |
|---|---|---|---|---|---|---|---|---|---|---|---|
| | $\xi$ | | | | | | | | | | |
| | 0 | 1 | 2 | 3 | 4 | 5 | 6 | 7 | 8 | 9 | 10 |

| | | | | | | | | | | | |
|---|---|---|---|---|---|---|---|---|---|---|---|
| Inner | 57.5 | 55.7 | 55.2 | 55.2 | 55.2 | 55.2 | 55.2 | 55.2 | 54.0 | 54.0 | 54.0 |
| Outer | 0.0 | 0.0 | 0.0 | 0.0 | 0.0 | 5.6 | 5.6 | 5.6 | 0.0 | 0.0 | 0.0 |
| Cytoplasm | 80.8 | 82.2 | 83.2 | 85.6 | 84.6 | 84.1 | 85.1 | 85.1 | 84.1 | 84.1 | 84.1 |
| Extracellular | 60.2 | 56.9 | 57.7 | 59.3 | 54.5 | 55.3 | 53.7 | 54.5 | 54.5 | 54.5 | 54.5 |
| OLA | 65.4 | 64.6 | 65.0 | 66.3 | 64.8 | 65.0 | 65.0 | 65.2 | 64.2 | 64.2 | 64.2 |
| OAA | 64.7 | 63.8 | 64.5 | 65.9 | 64.4 | 64.5 | 63.8 | 64.4 | 63.4 | 63.4 | 63.2 |

**Table S7**

Select different parameter $L$ to obtain prediction results of the Gram-negative dataset.

| Locations | Jackknife method (%) | | | | | | | | |
|---|---|---|---|---|---|---|---|---|---|
| | $L$ | | | | | | | | |
| | 5 | 10 | 15 | 20 | 25 | 30 | 35 | 40 | 45 |
| Cell inner membrane | 65.9 | 65.7 | 66.8 | 65.9 | 66.2 | 66.6 | 66.8 | 66.4 | 67.1 |
| Cell outer membrane | 0.0 | 0.0 | 0.0 | 0.0 | 0.8 | 0.0 | 0.8 | 0.0 | 0.0 |
| Cytoplasm | 62.2 | 62.7 | 63.7 | 65.6 | 63.2 | 65.1 | 66.3 | 67.6 | 66.8 |
| Extracellular | 4.5 | 6.0 | 8.3 | 3.0 | 7.5 | 9.0 | 8.3 | 6.8 | 9.0 |
| Fimbrium | 0.0 | 0.0 | 9.4 | 3.1 | 0.0 | 0.0 | 0.0 | 0.0 | 0.0 |
| Flagellum | 0.0 | 0.0 | 0.0 | 0.0 | 0.0 | 0.0 | 8.3 | 8.3 | 8.3 |
| Nucleoid | 0.0 | 0.0 | 0.0 | 0.0 | 0.0 | 0.0 | 0.0 | 0.0 | 0.0 |
| Periplasm | 0.0 | 0.6 | 1.1 | 2.8 | 0.6 | 1.7 | 2.8 | 3.3 | 3.3 |
| OLA | 43.1 | 43.4 | 44.6 | 44.4 | 44.0 | 44.8 | 45.5 | 45.5 | 45.8 |
| OAA | 43.0 | 43.3 | 44.3 | 44.2 | 43.7 | 44.4 | 44.9 | 45.3 | 45.4 |

**Table S8**

Select different parameter $L$ to obtain prediction results of the Gram-positive dataset.

| Locations | Jackknife method (%) | | | | | | | | |
|---|---|---|---|---|---|---|---|---|---|
| | $L$ | | | | | | | | |
| | 5 | 10 | 15 | 20 | 25 | 30 | 35 | 40 | 45 |
| Cell inner membrane | 48.3 | 48.3 | 47.7 | 47.7 | 48.3 | 47.7 | 47.7 | 47.7 | 47.7 |
| Cell outer membrane | 0.0 | 0.0 | 0.0 | 0.0 | 0.0 | 0.0 | 0.0 | 0.0 | 0.0 |
| Cytoplasm | 72.6 | 73.1 | 74.5 | 75.0 | 75.5 | 75.5 | 76.9 | 76.9 | 75.5 |
| Extracellular | 27.6 | 31.7 | 26.8 | 30.9 | 30.9 | 28.5 | 30.9 | 31.7 | 30.9 |
| OLA | 51.4 | 52.6 | 51.8 | 53.0 | 53.3 | 52.6 | 53.7 | 53.9 | 53.2 |
| OAA | 51.4 | 52.4 | 51.8 | 53.2 | 53.2 | 52.6 | 53.6 | 53.9 | 53.4 |

**Table S9**

Predicting the results of different feature extraction methods for Gram-positive and Gram-negative bacteria data sets.

| Jackknife method (%) |
|---|

| Datasets | Locations | Algorithms | | | | | |
|---|---|---|---|---|---|---|---|
| | | PseAAC | PsePSSM | EBGW | DC | GO | ALL |
| Gram-negative | Cell inner membrane | 75.9 | 75.6 | 66.4 | 77.6 | 95.9 | 99.5 |
| | Cell outer membrane | 33.1 | 37.1 | 0.0 | 31.5 | 86.3 | 96.8 |
| | Cytoplasm | 71.5 | 76.3 | 67.6 | 59.8 | 93.2 | 98.3 |
| | Extracellular | 16.5 | 41.4 | 6.8 | 18.8 | 90.2 | 98.5 |
| | Fimbrium | 0.0 | 31.3 | 0.0 | 21.9 | 78.1 | 96.9 |
| | Flagellum | 8.3 | 91.7 | 8.3 | 25.0 | 91.7 | 75.0 |
| | Nucleoid | 0.0 | 0.0 | 0.0 | 0.0 | 37.5 | 100 |
| | Periplasm | 61.1 | 50.0 | 3.3 | 17.2 | 93.9 | 99.4 |
| | OAA | 60.6 | 62.7 | 45.3 | 52.7 | 90.9 | **98.6** |
| | OLA | 61.1 | 65.0 | 45.5 | 53.7 | 92.8 | **98.6** |
| Gram-positive | Inner | 50.0 | 55.2 | 47.7 | 54.0 | 94.8 | 99.4 |
| | Outer | 5.6 | 0.0 | 0.0 | 11.1 | 44.4 | 94.4 |
| | Cytoplasm | 73.6 | 85.6 | 76.9 | 63.9 | 96.2 | 100 |
| | Extracellular | 48.8 | 59.3 | 31.7 | 41.5 | 89.4 | 99.2 |
| | OAA | 56.8 | 65.9 | 53.9 | 52.4 | 89.6 | **99.6** |
| | OLA | 57.6 | 66.3 | 53.9 | 53.5 | 92.4 | **99.4** |

ALL：PseAAC+PsePSSM+Dipeptide Composition+GO (DMLDA)

**Table S10**

Selecting different dimensionality reduction methods and different dimensions to obtain the overall actual accuracy results of subcellular localization.

| Datasets | Algorithms | Jackknife method (%) | | | | | | | | | |
|---|---|---|---|---|---|---|---|---|---|---|---|
| | | Dimensions | | | | | | | | | |
| | | 10 | 20 | 30 | 40 | 50 | 60 | 70 | 80 | 90 | 100 |
| Gram-negative | PCA | 61.8 | 67.7 | 66.7 | 67.2 | 67.3 | 68.3 | 68.5 | 68.6 | **70.0** | **70.0** |
| | MLSI | 67.3 | 73.1 | 72.8 | 73.6 | 74.1 | 73.9 | 74.9 | 74.8 | 75.1 | **75.8** |
| | MDDM | 82.8 | **83.5** | 82.9 | 82.9 | 82.2 | 81.9 | 82.3 | 81.0 | 80.7 | 80.9 |
| | MVMD | **76.1** | 73.3 | 73.2 | 72.6 | 72.1 | 72.2 | 71.0 | 71.2 | 72.4 | 72.9 |
| | DMLDA | **98.7** | 98.6 | 98.3 | 98.6 | 98.6 | 98.6 | 98.3 | 98.0 | 97.9 | 97.4 |
| Gram-positive | PCA | 64.7 | 63.2 | 66.7 | 65.3 | 67.8 | 66.3 | 68.2 | **69.2** | 68.8 | 68.8 |
| | MLSI | 84.6 | 85.2 | 84.4 | 85.4 | 85.5 | 86.3 | 85.2 | **87.1** | 85.0 | 84.6 |
| | MDDM | 82.8 | **83.5** | 82.9 | 82.9 | 82.2 | 81.9 | 82.3 | 81.0 | 80.7 | 80.9 |
| | MVMD | **75.0** | 72.4 | 70.1 | 70.3 | 69.2 | 71.1 | 68.0 | 69.2 | 72.1 | 69.7 |
| | DMLDA | 99.2 | 99.2 | 99.4 | 99.4 | 99.4 | **99.6** | 99.2 | 99.2 | 99.2 | 99.2 |

**Table S11**

Prediction results of Gram-negative dataset protein subcellular localization in different classification algorithms.

| Locations | Jackknife method (%) | | | |
|---|---|---|---|---|
| | Classifiers | | | |
| | MLkNN | ML-LOC | INSDIF | LIFT |
| Cell inner membrane | 99.5 | 99.5 | 99.5 | 99.5 |
| Cell outer membrane | 92.7 | 97.6 | 100 | 96.8 |
| Cytoplasm | 100 | 97.6 | 99.8 | 98.3 |
| Extracellular | 100 | 100 | 99.2 | 98.5 |
| Fimbrium | 96.9 | 96.9 | 100 | 96.9 |
| Flagellum | 100 | 0.0 | 0.0 | 75.0 |
| Nucleoid | 100 | 87.5 | 100 | 100 |
| Periplasm | 99.4 | 98.3 | 99.4 | 99.4 |
| OLA | 99.0 | 97.7 | 98.8 | 98.6 |
| OAA | 96.9 | 97.7 | 98.3 | 98.6 |

**Table S12**

Prediction results of Gram-positive dataset protein subcellular localization in different classification algorithms.

| Locations | Jackknife method (%) | | | |
|---|---|---|---|---|
| | Classifiers | | | |
| | MLkNN | ML-LOC | INSDIF | LIFT |
| Cell inner membrane | 100 | 98.9 | 100 | 99.4 |
| Cell outer membrane | 94.4 | 94.4 | 88.9 | 94.4 |
| Cytoplasm | 99.5 | 100 | 100 | 100 |
| Extracellular | 99.2 | 99.2 | 99.2 | 99.2 |
| OLA | 99.4 | 99.2 | 99.4 | 99.4 |
| OAA | 99.2 | 99.4 | 99.4 | 99.6 |

### 2. Supplementary Figure

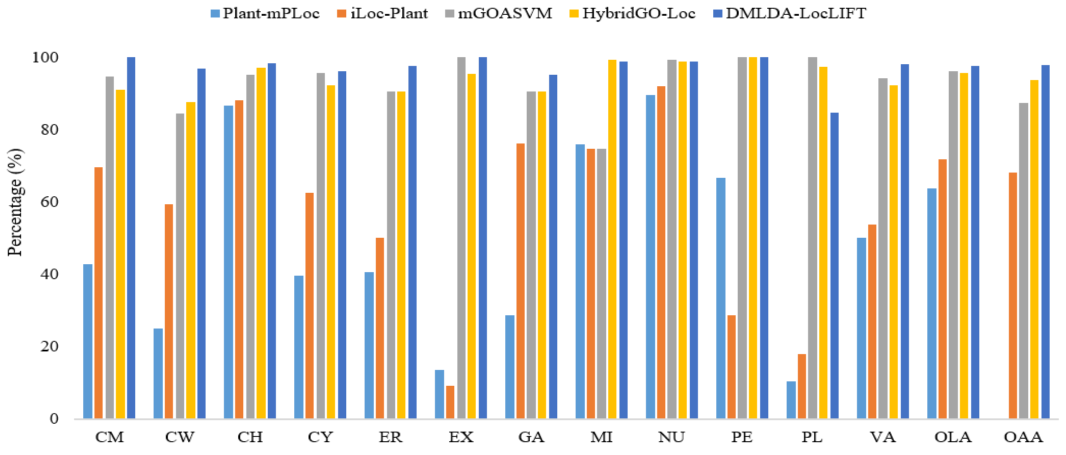

**Fig. S1.** Comparison of different prediction methods for protein subcellular localization on plant dataset. CM: Cell membrane; CW: Cell wall; CH: Chloroplast; CY: Cytoplasm; ER: Endoplasmic reticulum; EX: Extracellular; GA: Golgi apparatus; MI: Mitochondrion; NU: Nucleus; PE: Peroxisome; PL: Plastid; VA: Vacuole; OLA: overall location accuracy; OAA: overall actual accuracy. Plant-mPLoc does not provide the results of the overall actual accuracy
